# Formate Promotes Invasion and Metastasis by Activating Fatty Acid Synthesis and Matrix Metalloproteinases

**DOI:** 10.1101/2023.01.23.525172

**Authors:** Catherine Delbrouck, Nicole Kiweler, Vitaly I. Pozdeev, Laura Neises, Anaïs Oudin, Anne Schuster, Aymeric Fouquier d’Hérouël, Ruolin Shen, Rashi Halder, Antoine Lesur, Christoph Ogris, Nadia I. Lorenz, Christian Jaeger, Michael W. Ronellenfitsch, Marie Piraud, Alexander Skupin, Simone P. Niclou, Elisabeth Letellier, Johannes Meiser

## Abstract

Metabolic rewiring is essential to enable cancer onset and progression. One important metabolic pathway that is often hijacked by cancer cells is the one-carbon cycle, in which the third carbon of serine is oxidized to formate. We have previously shown that formate production in cancer cells often exceeds the anabolic demand, resulting in formate overflow. Furthermore, we observed that high extracellular formate promotes the *in vitro* invasiveness of glioblastoma (GBM) cells. However, additional data supporting this *in vitro* observation and mechanistic details remained elusive so far.

In the present study, we now demonstrate that inhibition of formate overflow results in a decreased invasiveness of GBM cells *ex vivo* and *in vivo*. Additionally, we observed that exposure to exogeneous formate can induce a transiently stable pro-invasive phenotype that results in increased metastasis formation *in vivo*. All in all, these results suggest that a local formate increase within the tumor microenvironment may be one factor that can promote cancer cell motility and dissemination.

Mechanistically, we uncover a previously undescribed interplay where formate acts as a trigger to alter fatty acid metabolism and matrix metalloproteinase (MMP) activity which in turn impacts cancer cell invasiveness. We thus highlight the role of formate as a pro-invasive metabolite. Gaining a deeper understanding of formate overflow and how it promotes invasion in cancer, may open new therapeutic opportunities to prevent cancer cell dissmination.

## INTRODUCTION

The spreading of cancer cells is a complex and sequential process, initiated by migration and local invasion from the primary tumor site into the surrounding tissue. Cell invasion requires distinct metabolic reprogramming to meet the cell’s specific needs. Proliferating cancer cells display a metabolism that is adapted towards an anabolic program to generate metabolic intermediates required for cell division. In contrast, invasive disseminating cells rewire their metabolism towards a catabolic program to support the relative increase in RedOx and energy demand (Benzarti et al., 2020; Bergers and Fendt, 2021; Elia et al., 2018).

One-Carbon (1C) metabolism supports the synthesis of different metabolites that serve as building blocks of the cell, including purines. In the first step of the cycle, carbon-3 of serine is transferred to tetrahydrofolate (THF) and is oxidised subsequently through the cycle (Brosnan and Brosnan, 2016; Tibbetts and Appling, 2010). The 1C metabolism does not only support biosynthesis, but has also relevance in the context of RedOx control, bioenergetics and methylation. The plasticity of the pathway allows cancer cells to exploit it according to their current cellular needs. Within the mitochondrion, serine is catabolized to glycine and formate yielding reducing equivalents and ATP. In the cytosolic part of the 1C cycle, both glycine and formate can subsequently serve as precursors for nucleotides, proteins or glutathione (GSH) (Benzarti et al., 2020; Young et al., 2017) or is used to regenerate serine, closing the cycle. Yet, even in fully proliferating cells not all formate is used for nucleotide synthesis and excessive amounts of formate are released from cancer cells resulting in a formate overflow phenotype (Meiser et al., 2016). Methylenetetrahydrofolate dehydrogenase 1 like (MTHFD1L) is an ATP producing enzyme of the mitochondrial 1C metabolism involved in formate generation and essential for formate overflow (Christensen et al., 2005; Meiser et al., 2016). While the dependency of formate overflow on active mitochondrial 1C metabolism has been demonstrated (Kiweler et al., 2022; Meiser et al., 2018; Meiser et al., 2016), only little is known about its selective advantage in cancer cells.

Recent *in vitro* analyses in our laboratory revealed that formate overflow promotes invasion of malignant brain tumor (Glioblastoma (GBM)) cells *in vitro*. The pro-invasive effect upon formate was concentration-dependent with significant changes observed starting form 100 μM, suggesting that high formate concentrations may serve as an auto- or paracrine signal to increase the invasiveness of GBM cells (Meiser et al., 2018). Yet, the physiological relevance and mechanistic details of this observation remain to be addressed.

In the present study, we employed *ex vivo* and *in vivo* approaches and provide evidence that the formate-dependent effect on GBM invasion is physiologically relevant. We recapitulate our findings in various breast cancer models and show that formate does not only impact local invasion of cancer cells but also the onset of breast cancer metastasis *in vivo*. Mechanistically, we uncover that formate promotes fatty acid synthesis and the excretion of matrix metalloproteases (MMPs). Our data suggest a previously undescribed interplay between 1C metabolism and lipid metabolism where formate might act as a trigger to alter fatty acid metabolism which in turn impacts cancer cell invasiveness and the metastatic potential. Targeting either, MMP release or fatty acid synthesis *in vitro* and *ex vivo*, suppresses the formate-dependent pro-invasive effect.

We postulate that targeting formate overflow in cancer cells to reduce the formate concentration in the tumor microenvironment may represent an attractive approach to limit cancer cell dissemination.

## RESULTS

### Formate specifically induces glioblastoma cell invasion

To extend our previous findings (Meiser et al., 2018), we investigated the effect of formate on the invasion capacity of a panel of patient-derived GBM cell lines including BG5, BG7, GG6 and GG16 (Figure 1A; Supplementary Figure 1A; Supplementary Figure 1B). Although basic invasion efficiency is cell line dependent (Schuster et al., 2020), all tested cell lines showed increased invasion after formate treatment.

**Figure 1:**
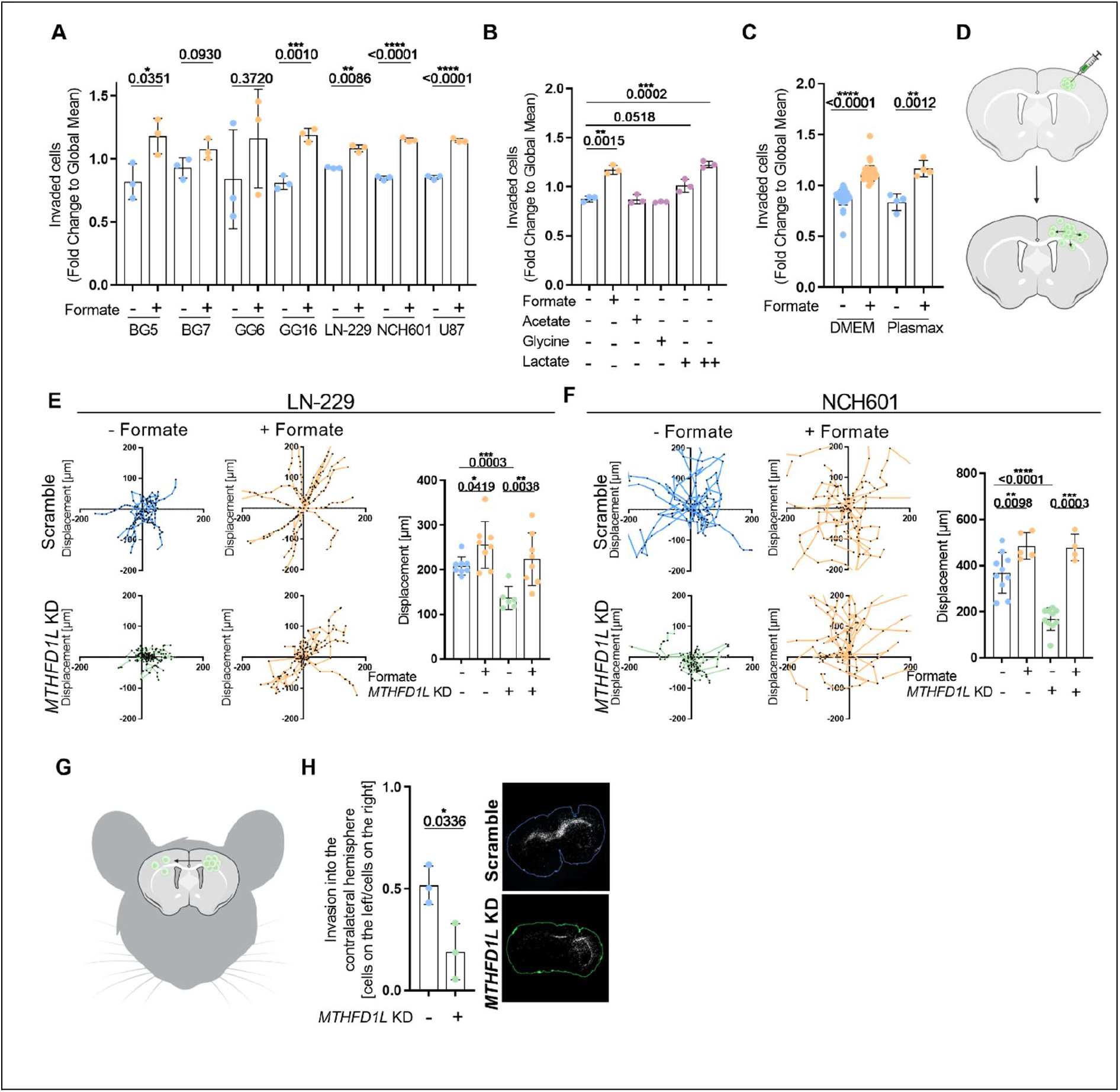
Formate Is a Specific Activator of Glioblastoma Cell Invasion. **(A)** Invasion of different glioblastoma cells: BG5, BG7, GG6, GG16, LN-229, NCH601 and U87 was assessed using ECM-Collagen coated Boyden Chambers. The cells were treated or not with 500 μM formate. Each dot represents an independent experiment; mean ± SD; Unpaired *t*-test with Welch’s correction. **(B)** Invasion of LN-229 cells which were treated with 500 μM Na-formate; 500 μM Na-acetate; 500 μM glycine; 500 μM Na-lactate or 2 mM Na-lactate was assessed using ECM-Collagen coated Boyden chambers. Each dot represents an independent experiment; mean ± SD; Unpaired *t*-test with Welch’s correction. **(C)** Comparison of the invasive phenotype of LN-229 cells which were treated for 24 hours with 500 μM formate and cultured in different medium (Dulbecco’s modified Eagle’s medium (DMEM) or selfmade physiological medium (Plasmax)). Invasion was assessed using ECM-Collagen coated Boyden Chambers. Each dot represents an independent experiment; mean ± SD; Unpaired *t*-test with Welch’s correction. **(D)** Shematic representation of the *ex vivo* brain slice assay. **(E)** Displacement of LN-229 (*sh*Scramble vs. *MTHFD1L* KD) cells injected in *ex vivo* brain slice cultures. The cells were cultured during the entire experiment with 500 μM Na-formate. Results are displayed as mean ± SEM; Unpaired *t*-test with Welch’s correction. Each dot represents the average displacement of 15 cells. **(F)** Displacement of NCH601 (*sh*Scramble vs. *MTHFD1L* KD) cells injected in *ex vivo* brain slice cultures. The cells were cultured during the entire experiment with 500 μM Na-formate. Results are displayed as mean ± SEM; Unpaired *t*-test with Welch’s correction. Each dot represents the average displacement of 15 cells. **(G)** Shematic representation of the *in vivo* brain tumor model **(H)** Invasion of NCH601 (*sh*Scramble *vs*. *MTHFD1L* KD) cells into the contralateral hemisphere of NU/NU Nude female mice after 4 months. Each dot represents an individual mouse; mean ± SD; Unpaired *t*-test with Welch’s correction.

To investigate if the observed pro-invasive effect is specific to formate, we performed *in vitro* invasion assays with different small metabolites to compare their impact on the invasiveness of the LN-229 cell line (Figure 1B; Supplementary Figure 1D). Acetate, which differs from formate only by one additional methyl group, had no effect on cell invasion (Figure 1B). Glycine which, in case of serine catabolism, is often released in an 1:1 ratio with formate (Meiser et al., 2016), also had no impact on the invasiveness of LN-229 cells (Figure 1B). Lactate, known to promote invasion (Sonveaux et al., 2008), served as a positive control and promoted the invasion rate of LN-229 cells in a dose-dependent manner (Figure 1B). Of note, the promoting effect on invasion at 2 mM of Na-lactate was in a similar range as observed upon addition of 500 μM of Na-formate. In line with these results, we did not observe a growth reduction upon these different treatments (Supplementary Figure 1C). In brief, these results consolidate that formate has a specific role in promoting cancer cell invasion.

To test the physiological relevance of the observed pro-invasive phenoytpe, we chose three different approaches. First, we performed *in vitro* Boyden chamber assays in Plasmax medium, a medium known to closely recapitulate the nutrient composition of human blood (Vande Voorde et al., 2019) (Figure 1C; Supplementary Figure 1E). Here, a comparable increase in invasiveness was detectable after the addition of 500 μM Na-formate. Second, to better mimic invasion in the brain microenvironment, we employed *ex vivo* brain slice assays. Mice brains were cut into 400 μm slices and transferred into a specific medium that allows culturing of viable brain tissue slides over several days. By using this model, cancer cells were injected into the brain slices to monitor their invasive capacity *ex vivo* at a single cell level (Figure 1D) (Eisemann et al., 2018; Schuster et al., 2020). To validate the decreased invasiveness upon *MTHFD1L* KD *ex vivo*, we implanted LN-229 cells or the highly invasive glioblastoma cell line NCH601 with sh*Ctrl* or sh*MTHFD1L* into brain slices cultured with or without 500 μM Na-formate (Supplementary Figure 1F-G). Suppression of formate overflow upon this KD has already been demonstrated (Meiser et al., 2018). In line with our reported *in vitro* data, the addition of 500 μM Na-formate significantly increased the single cell velocity of sh*Ctrl* cells. In contrast, stable sh*MTHFD1L* KD resulted in a significant reduction of cellular velocity relative to sh*Ctrl* cells. Finally, addition of exogenous formate rescued the reduced velocity of sh*MTHFD1L* cells (Figure 1E-F). Of note, neither sh*MTHFD1L* nor addition of 500 μM Na-formate altered the proliferation or the viability of LN-229 or NCH601 cells (Meiser et al., 2018). Third, we injected the cells intracranially into the right brain hemisphere of Nude mice to measure invasion into the contralateral hemisphere (Figure 1G). Two months after intracranial tumor implantation, no significant differences between the sh*Ctrl* and sh*MTHFD1L* cells were noticed (Supplementary Figure 1H), indicating similar viability and growth rates of sh*MTHFD1L* cells compared to sh*Ctrl* cells *in vivo*. However, four months after implantation, we observed a significantly lower rate of invasion into the contralateral hemisphere of sh*MTHFD1L* KD cells compared to sh*Ctrl* cells (Figure 1H).

Taken together, these data underline that increased extracellular formate concentrations promote cancer cell invasion *in vitro*, *ex vivo* and *in vivo* and that inhibition of formate overflow reduces the invasiveness in GBM *ex vivo* and *in vivo*.

### Formate primes cancer cells towards an invasive phenotype and promotes the seeding capacity of breast cancer cells *in vivo*

After consolidating that formate acts as a pro-invasive metabolite, we wanted to investigate if formate promotes not only local invasion but also influences the metastatic seeding capacity. As malignant gliomas only rarely metastasize from the central nervous system to systemic sites (Liwnicz and Rubinstein, 1979; Pasquier et al., 1980), we decided to study the role of formate also in context of breast cancer. First, to verify if formate also promotes invasiveness of breast cancer cells, we performed *in vitro* Boyden chamber assays with different breast cancer cell lines (BT20, 4T1, MDA-MB-231 and MDA-MB-468) (Figure 2A). Formate increased the invasive potential of all four tested cell lines.

**Figure 2.**
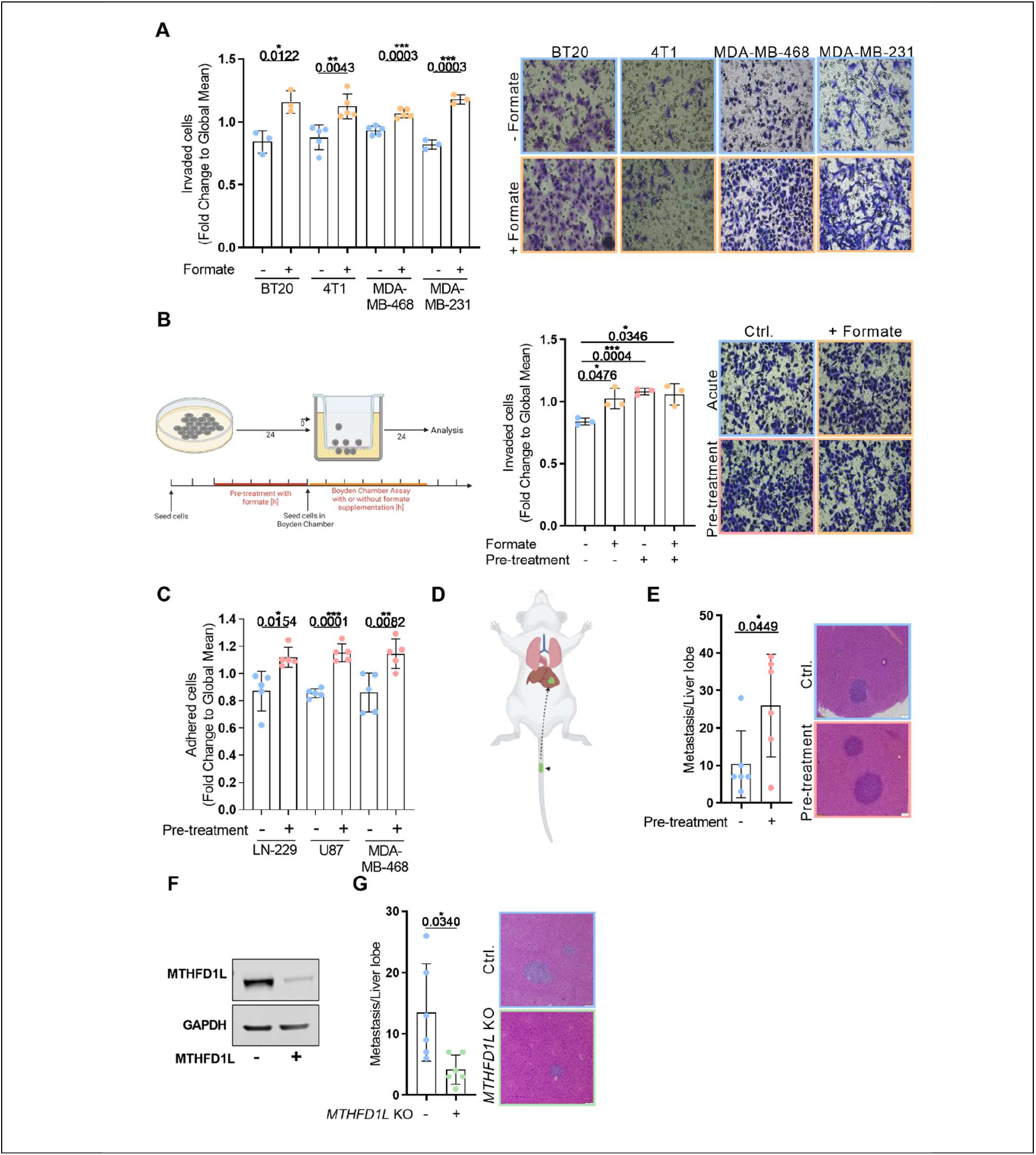
Formate Primes Cancer Cells towards an Invasive Phenotype and Promotes the Seeding Capacity of Breast Cancer Cells *in vivo*. **(A)** On the left: Invasion of different breast cancer cells: BT20, 4T1, MDA-MB-468 and MDA-MB-231 treated for 24 hours with 500 μM Na-formate was assed using ECM-Collagen coated Boyden chambers. Each dot represents an independent experiment; mean ± SD; Unpaired *t*-test with Welch’s correction. On the right: Representative pictures of the Boyden chamber assay of different breast cancer cells (BT20, 4T1, MDA-MB-468, and MDA-MB-231) treated with Na-formate. **(B)** Left: Shematic representation of the experimental layout. Right: Invasion of MDA-MB-468 cells pre-treated for 24 hours with 500 μM formate was assed using ECM-Collagen coated Boyden chambers. Each dot represents an independent experiment; mean ± SD; Unpaired *t*-test with Welch’s correction. **(C)** Adhesion of LN-229, U87 or MDA-MB-468 cells, which were pre-treated for 24 hours with formate was assed on ECM-Collagen-coated cover slips. Each dot represents an independent experiment; mean ± SD; Unpaired *t*-test with Welch’s correction. **(D)** Shematic representation of the *in vivo* tail vein model **(E)** On the left: Macroscopic metastases count in livers after injecting MDA-MB-468 control or 24 hours formate (500 μM) pre-treated cells in the tail vein of NSG mouse. On the right: Representative H&E staining of liver metastasis. **(F)** Representative Western Blot showing MTHFD1L expression in MDA-MB-468 control and *MTHFD1L* KD cells. **(G)** On the left: Macroscopic metastases count in livers after injecting MDA-MB-468 control or *MTHFD1L* KD cells in the tail vein of NSG mouse. On the right: Representative H&E staining of liver metastasis.

To test if formate can also prime cancer cells to induce a longer lasting effect, we pretreated cells for 24 hours with formate prior to performing *in vitro* invasion assays (Figure 2B). We observed that formate pre-treated cells had a similar invasive potential as the acutely treated cells. Addition of formate to pre-treated cells did not further increase the rate of invasion. To study if formate also primes GBM cells (LN-229 and U87), and to investigate if shorter exposure times are already sufficient, we pre-treated these cells for 3, 24 and 72 hours with 500 μM Na-formate before examining their invasivness in Boyden chamber assays (Supplementary Figure 2A). A significant increase in invasion was observed starting from 3 hours of formate treatment. Shorter time points did not impact cancer cell invasion, suggesting that 3 hours of formate is a minimum time required to prime the cells (Supplementary Figure 2B). Next, the stability of the pro-invasive signal was investigated. To test the stability of the signal, cancer cells were pre-treated for 24 hours with formate. Then, the treatment was removed and the cells were cultured for different time spans without formate prior invasion analysis. Despite all cell specific differences, we observed a signal stabilisation ranging from 3 hours in U87 to less than 1 hour in MDA-MB-468 cells (Supplementary Figure 2C).

As the pro-invasive signal is fading out overt time, we hypothesised that formate may play an important role in the initial steps of invasion and tumour cell escape. Adhesion represents a crucial and early step in the colonisation process. Thus, we investigated if formate pre-treatment affects the capacity of cancer cells to adhere. In all the tested cell lines (LN-229, U87, and MDA-MB-468) formate pre-treated cells adhered more compared to the untreated control (Figure 3C). In sum, we observed that formate can prime cancer cells to a pro-metastatic phenotype promoting both invasive and adhesive capacites.

**Figure 3:**
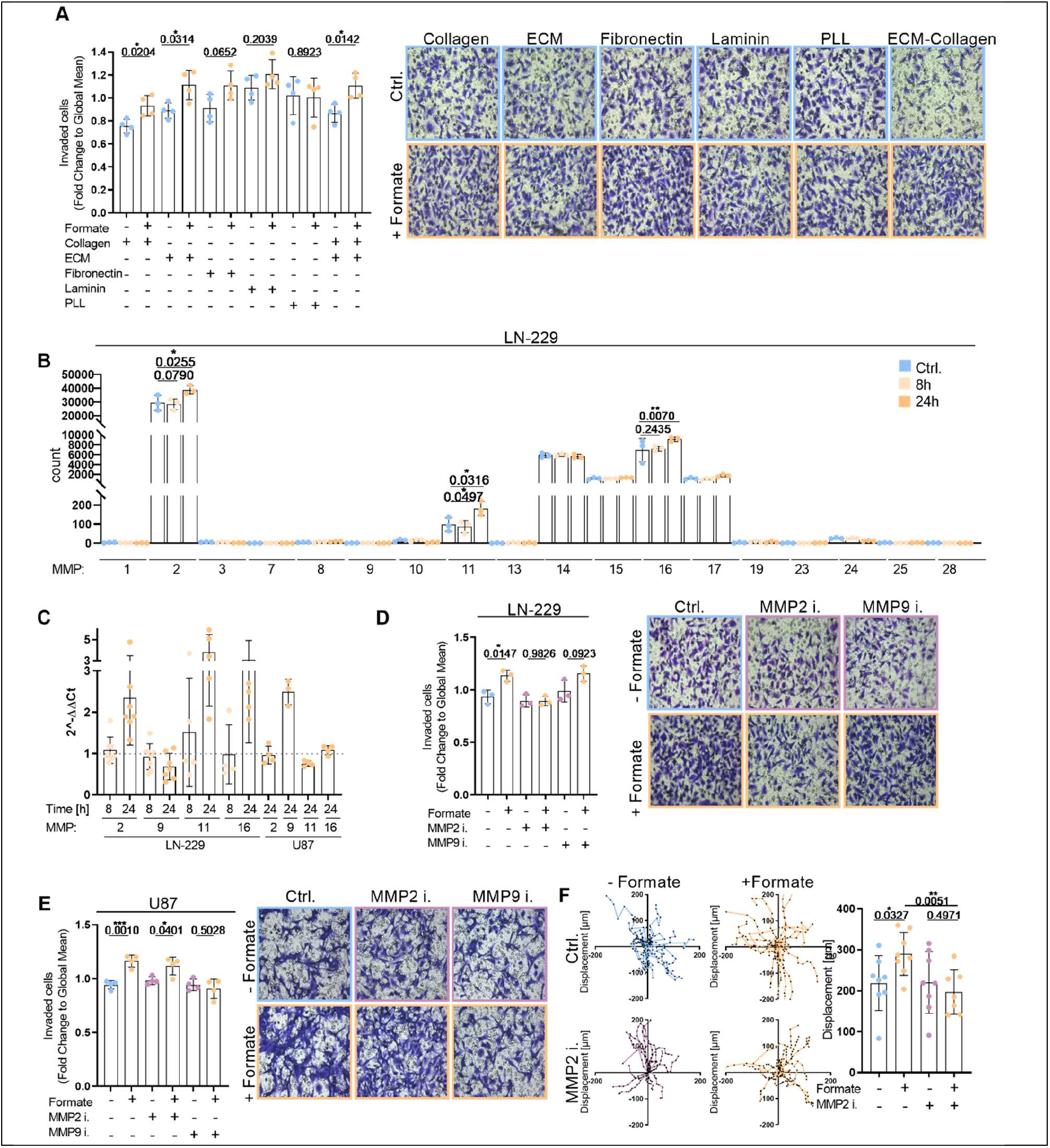
Formate Promotes Invasion in Dependence on Matrix Metalloproteinase. **(A)** Invasion of LN-229 cells treated for 24 hours with 500 μM Na-formate was assed on differently coated Boyden chambers (Collagen, ECM, Fibronectine, Laminin, PLL, and ECM-Collagen). Each dot represents an independent experiment; mean ± SD; Unpaired *t*-test with Welch’s correction. **(B)** RNAseq results displaying the count of different MMPs (MMP1, MMP2, MMP3, MMP7, MMP8, MMP9, MMP10, MMP11, MMP13, MMP14, MMP15, MMP16, MMP17, MMP19, MMP23, MMP24, MMP25, and MMP28) expression in LN-229 cells which were treated for 8 hours or 24 hours with 500 μM Na-formate. Each dot represents an independent experiment; mean ± SD; Unpaired *t*-test with Welch’s correction. mRNA expression of MMP2, MMP9, MMP11, and MMP16 in LN-229 and U87 cells which were treated for 8 h or 24 h with 500 μM Na-formate relative to untreated cells as measured using real-time RT-qPCR. Each dot represents an independent experiment; mean ± SD. **(D)** Invasion of LN-229 cells treated for 24 h with 500 μM formate and 5 μM MMP inhibitors (MMP2 inhibitor and MMP9 inhibitor) was assed using ECM-Collagen coated Boyden chambers. Each dot represents an independent experiment; mean ± SD; Unpaired *t*-test with Welch’s correction. **(E)** Invasion of U87 cells treated for 24 h with 500 μM formate and 5 μM MMP inhibitors (MMP2 inhibitor and MMP9 inhibitor) was assed using ECM-Collagen coated Boyden chambers. Each dot represents an independent experiment; mean ± SD; Unpaired *t*-test with Welch’s correction. **(F)** Displacement of LN-229 cells injected in *ex vivo* brain slices. The cells were cultured during the entire experiment with 500 μM formate, 5 μM ARP100, or the combination of both. Results are displayed as mean ± SEM; Unpaired *t*-test with Welch’s correction. Each dot represent the average displacement of 15 cells.

To test if such formate-dependent priming impacts metastasis formation, we investigated the *in vivo* seeding capacity of formate pretreated MDA-MB-468 cells (Figure 2D). To do so, we injected formate pretreated MDA-MB-468 cells into the tail vein of mice. Six weeks after injection, we found a significantly higher rate of liver metastases in mice that were injected with the formate pretreated cells (Figure 2E). Subsequently, we injected *MTHFD1L* KO cells (Figure 2F) which are formate overflow negative (Kiweler et al., 2022). Here we noticed a significantly lower rate of liver metastases compared to the Ctrl cells (Figure 2G), which is in line with previous work of our laboratory using an orthotopic breast cancer model (Kiweler et al., 2022).

In summary, we conclude that formate not only promotes local invasion but also promotes the seeding capacity of breast cancer cells *in vivo* and that by preventing mitochondrial formate production, invasion and metastasis can be reduced.

**Formate promotes invasion by activation of matrix metalloproteinases**

In addition to increased capacity to adhere, the proteolysis of the extracellular-matrix (ECM)(Liotta, 2016) is also an important process contributing to metastasis. To reveal by which means high formate levels promote cancer cell invasion, we tried to investigate separately which of these two features are influenced by formate. To evaluate the migratory potential of formate treated cells, we performed Boyden chamber (Supplementary Figure 3A) and wound healing (scratch) assays (Supplementary Figure 3B) without ECM-collagen coating. Both assays revealed that formate did not further increase the migratory potential of different analyzed cell lines (Supplementary Figure 3A-B). Next, we investigated the capacity of cancer cells to degrade the extracellular matrix (ECM). To that end, we explored if the type of coating impacts the observed effect of formate on invasion. Thus, we compared the invasion capacity in respect to six different coatings (Figure 3A). We noticed that the formate-dependent pro-invasive effect was strongly impacted by the type of coating. Cells that had to invade through collagen, ECM or a collagen-ECM mix showed increased invasiveness upon addition of formate. Cells invading through fibronectin and laminin showed a trend of increased invasiveness upon formate. However, poly-L-lysine (PLL) coating did not result in increased transfer of LN-229 cells. As PLL coating do not require active matrix proteolysis by the cells, resulting in a rather migration like assay, we speculate that formate has no effect in these kind of matrices. This assumption is in line with our previous observation that formate does not further increase the cell’s migratory capacity (Supplementary Figure 3A-B). In contrast, the positive effect observed on collagen and ECM matrices is in favour of our hypothesis that formate increases the cellular capacity to degrade the matrix. To test this hypothesis, we profiled the abundance of all detectable matrix metalloproteinases (MMPs) by RNA-seq in LN-229 cells (Figure 3B). LN-229 were treated for 8 or 24 hours with 500 μM of formate prior RNA extraction. Among the 18 MMPs that could be detected, three MMPs showed significant increased expression after 24 hours of formate treatment: MMP2, MMP11 and MMP16. Among these three MMPs, MMP2 showed by far the strongest overall expression level. MMP2 builds together with MMP9 the distinct subclass of gelatinases within the MMP family. Furthermore, MMP2 and MMP9, have been described as contributors to the progression of malignant glioma (Fabian et al., 2021; Guo et al., 2005; Nakada et al., 2003; Sincevičiūtė et al., 2018; Vehlow and Cordes, 2013). Therefore, we focused our subsequent analyses on MMP2 and MMP9.

Thus, we first confirmed the obtained RNAseq-data by RT-qPCR. In line with the RNAseq analysis, MMP2, MMP11 and MMP16 gene expression in LN-229 cells was increased after 24 h of formate treatment while MMP9 gene expression remained unchanged (Figure 3C). Interestingly, U87 cells showed increased gene expression of MMP9 only (Figure 3C).The breast cancer models (MDA-MB-468, and MDA-MB-231) showed only weak induction of MMP’s (Supplementary Figure 3C). In conclusion, we observed a formate-dependent activation of MMPs in particular in glioblastoma models, yet the specific MMP that is activated upon formate is cell line dependent.

To test the functional relevance the importance of MMPs in formate-induced invasion, we measured the release of MMP2 and MMP9 protein by LN-229 cells into the cell culture medium after 500 μM Na-formate treatment (Supplementary Figure 3D). MMP9 release was detected across all conditions, but no significant changes upon formate treatment were measured. Interestingly, MMP2 release was coating-dependent, as MMP2 protein could only be detected in the medium when the cancer cells were cultured on an ECM-collagen coating. Following quantification of the protein signal, subtle changes have been observed between the conditions (Supplementary Figure 3D).

To further test induction of MMP2 in LN-229 cells upon formate treatment, we performed gelatine zymography assays to study MMP activity (Supplementary Figure 3E). Here, we identified increased MMP2 activity in formate treated cells. In contrast, stable sh*MTHFD1L* KD resulted in a significant reduction relative to the sh*Ctrl*. Finally, addition of exogenous formate partially rescued the reduced MMP2 activity of sh*MTHFD1L* cells. In line with our gene expression results, no changes of MMP9 activity were monitored (Supplementary Figure 3E).

Having identified a correlation between formate and MMPs, we investigated if formate-induced invasion is also functionally related to MMP abundance and activity. To achieve this, we performed invasion assays in combination with formate treatment and MMP inhibition (MMPi). In LN-229 cells, we observed that MMP2i (ARP-100) prevented formate-induced invasion while MMP9i (MMP-9 Inhibitor I) had no repressive effect on the formate-dependent increase of invasion (Figure 3D). These results were in line with our gene expression analysis. Consequently, in U87 cells we expected a specific effect caused by MMP9i and not with MMP2i. Indeed, in U87 cells only MMP9i but not MMP2i suppressed formate induced invasion (Figure 3E). Of note, the observed decreased invasiveness is not a result of a decreased proliferation rate (Supplemetary Figure 3F). To test the physiological relevance of this mechanism, we performed *ex vivo* brain slice assays. To that end, we implanted LN-229 cells and tested the formate-dependent effect on invasion in respect to MMP2 inhibition (Figure 3F). In line with previously shown data (Figure 1D), addition of 500 μM of Na-formate resulted in a significant increase in single cell velocity of sh*Ctrl* cells. In contrast, MMP2i prevented the formate-induced increase in the cellular movement, indicating that MMP2i can block the formate-dependent pro-invasive effect.

In summary, these data indicate that formate promotes invasion via activation of matrix metalloproteinases.

### Targeting fatty acid synthesis inhibits formate-dependent invasion

Next, we aimed to identify additional formate-dependent factors that operate upstream of MMPs by following both knowledge-guided hypothesis testing and unbiased profiling using metabolomics and transcriptomics.

It has been demonstrated before that mitochondrial ROS promotes cancer cell invasion (Cheung et al., 2020; Porporato et al., 2014) and that formate can increase mitochondrial ROS production in murine liver mitochondria (Young et al., 2017). Different ROS scavengers were successful in preventing a formate-dependent increase in invasion without affecting the proliferation rate (Supplementary Figure 4A), suggesting that ROS might be one factor that is involved in the signaling cascade. However, we were not able to detect differences in ROS levels despite different technical attempts. To better indentify factors involved in the regulatory network, we employed additional unbiased approaches.

**Figure 4:**
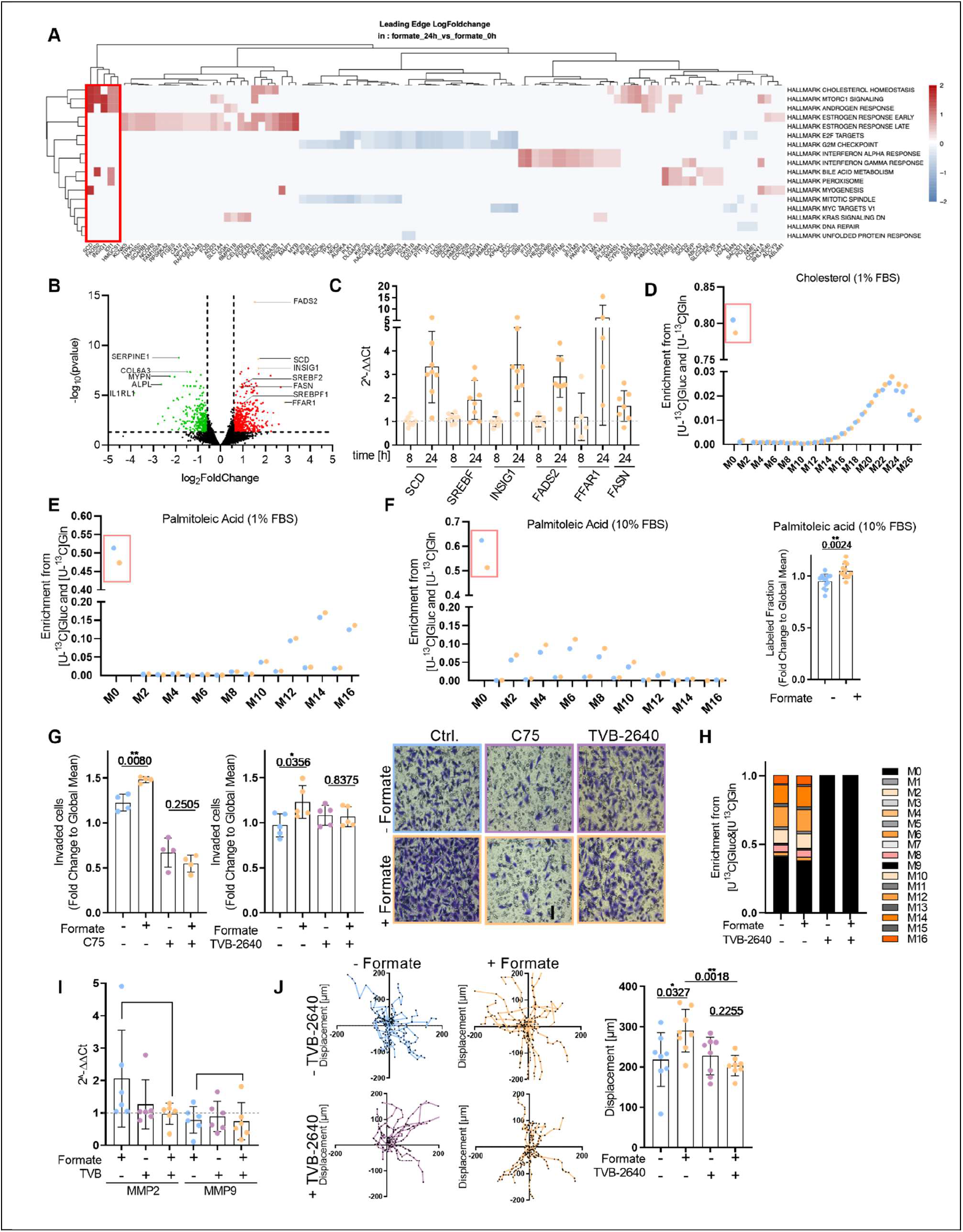
Targeting Fatty Acid Synthesis Inhibits Formate Dependent Invasion. **(A)** Gene set enrichment analysis (GSEA) representing LN-229 cells treated for 24 h with 500 μM formate relative to LN-229 control cells as measured using RNAseq data. **(B)** Volcano Blot representing LN-229 cells treated for 24 h with 500 μM formate relative to LN-229 control cells as measured using RNAseq data. **(C)** mRNA expression from SCD, SREBF, INSIG1, FADS2, FFAR1 and FASN in LN-229 cells which were treated for 8 h or 24 h with 500 μM formate relative to non-treated LN-229 cells as measured using real-time RT-qPCR. Each dot represents an independent experiment. Representative graph of isotopologues enrichement of Cholesterol in DMEM supplemeneted with 1% FBS upon [U-^13^C]glucose and [U-^13^C]glutamine tracer response to 72 h 500 μM formate treatment in LN-229 cells. Experiment was repeated 3 times with triplicate wells. **(E)** Representative graph of isotopologues enrichement of Hexadecenoic acid 2 in DMEM supplemeneted with 1% FBS upon [U-^13^C]glucose and [U-^13^C]glutamine tracer in response to 72 h 500 μM formate treatment in LN-229 cells. **(F)** On the left: Representative graph of isotopologues enrichement of Hexadecenoic acid 2 in DMEM supplemented with 10% FBS upon [U-^13^C]glucose and [U-^13^C]glutamine tracer in response to 72 h 500 μM formate treatment in LN-229 cells. Experiment was repeated 3 times with triplicate wells. On the right: Labeled fraction of Hexadecenoic acid 2 upon [U-^13^C]glucose and [U-^13^C]glutamine tracer in response to 72 h 500 μM formate treatment in LN-229 cells. Each dot represents an independent experiment with triplicate wells; mean ± SD; Unpaired *t*-test with Welch’s correction. **(G)** Invasion of LN-229 cells treated for 24 h with 500 μM formate and a FASN inhibitor (10 μM C75 and 10 μM TVB-2640) was assed using ECM-collagen coated Boyden chambers. Each dot represents an independent experiment; mean ± SD; Unpaired *t*-test with Welch’s correction. **(H)** Representative graph of isotopologue enrichment of hexadecenoic acid after [U-^13^C]glucose and [U-^13^C]glutamine tracer supplementation in response to 72 hours of 500 μM formate and 10 μM TVB-2640 treatment in LN-229 cells. **(I)** mRNA expression from MMP2 and MMP9 in LN-229 cells which were treated for 24 hours with 500 μM formate and 10 μM TVB-2640 relative to LN-229 untreated control cells as measured using real-time RT-qPCR. Each dot represents an independent experiment; mean ± SD. **(J)** Displacement of LN-229 cells injected in *ex vivo* brain slices. The cells were cultured during the entire experiment with 500 μM formate, 10 μM TVB-2640, or the combination of both. Results are displayed as mean ± SEM; Unpaired *t*-test with Welch’s correction. Each dot represent the average displacement of 15 cells.

First, we investigated if increased formate concentrations cause global metabolic alterations of the central carbon metabolism. Unexpectedly, no changes were detected in metabolite steady-state levels, metabolic fluxes or in the cellular respiration rate (Supplementary Figure 4B-D).

Next, we analyzed the RNAseq dataset of formate-treated LN-229 cells in greater detail. While eight hours of formate treatment had no impact on the transcriptome (data not shown), 24 hours of treatment induced profound changes (Figure 4A-B & Supplementary Table 1). Gene set enrichment analysis (GSEA) pinpointed several specific subsets of the transcriptome that were affected by 24 hours formate treatment (Figure 4A). Among the different groups identified, we uncovered several up-reguated gene sets that shared a common biological function. Interestingly, “cholesterol homeostasis”, “androgen response”, “estrogen early response” and “estrogen late response” were the most differentially expressed gene sets. As androgens and estrogens are made from cholesterol all these gene sets route back to acetyl-CoA metabolism and the mevalonate pathway. Interestingly, mTORC1, which is a known upstream regulator of cholesterol biosynthesis (Laplante and Sabatini, 2009; Porstmann et al., 2009), was also identified as a top upregulated gene-set. As side note, a gene-set related to peroxisomes was also upregulated, which fits our observation related to ROS (Supplementary Figure 3A) and especially the potency of ROS scavengers to reduce a formate-dependent increase in cell invasion.

When considering the most strongly induced genes, many of those related to lipid metabolism: Fatty Acid Desaturase 2 (FADS2), Stearoyl-CoA desaturase-1 (SCD), Insulin-induced gene 1 (INSIG1), Free fatty acid receptor 1 (FFAR1), Fatty Acid Synthase (FASN) and Sterol regulatory element-binding protein 1 & 2 (SREBF1/2). Taken together, we observed that increased formate concentrations promote a characteristic gene signature that is related to a rewiring of lipid metabolism.

To validate this identified lipid signature, we performed RT-qPCR (Figure 4C & Supplementary Figure 4E). Accordingly, we found that formate increases the expression of several genes related to fatty acid synthesis (SCD, SREBF, INSIG1, FADS2, FFAR1 and FASN) in LN-229 (Figure 4C) and MDA-MB-468 cells (Supplementary Figure 4E), while U87 cells showed no activation of these genes (except FFAR1) (Supplementary Figure 4E).

To further validate the assumption that formate promotes fatty acid synthesis, we measured the protein level of the fatty acid synthase (FASN) in the LN-229 cells (Supplementary Figure 4F). However, upon formate we only observed subtle changes that varied across different time points, suggesting that not only protein level but also the actual metabolic rate (enzymatic activity) may be a relevant factor. To that end, we investigated the rate of fatty acid synthesis in a more direct approach and performed stable-isotope assisted metabolic flux analysis of cells that were treated with or without formate to monitor the rate of ^13^C incorporation into fatty acids and cholesterol (Figure 4E-F; Supplementary Figure 4G-I). To increase the labelled fraction of the acetyl-CoA pool and to avoid potential differences originating from different glucose and glutamine contribution to the acetyl-coA pool, we applied glucose and glutamine tracers simultaneously. Following this approach, we observed increased fatty acid synthesis upon formate (Figure 4F), but the overall rate of *de novo* cholesterol biosynthesis was neglectable (~4%) (Supplementary Figure 4G). To induce endogenous cholesterol synthesis we cultured the cells in 1% FBS. In this condition, we monitored an overall increased activity in cholesterol synthesis and that formate supplementation promoted cholesterol synthesis in LN-229 cells (Figure 4E). Of note, at 1% FBS, fatty acid synthesis remained increased upon formate.

As all the previous functional assays were performed in 10% FBS, we decided to test a functional relation between formate and fatty acid metabolism at 10% FBS. To that end, we performed *in vitro* invasion assays and targeted fatty acid synthesis. Two different fatty acid synthase (FASN) inhibitors, C75 and TVB-2640, successfully prevented the formate-dependent increase of invasion in LN-229 cells (Figure 4G) and MDA-MB-468 cells (Supplementary Figure 4K). C75 treatment resulted in a decreased rate of basal invasion. In contrast, TVB-2640 treatment. The latter inhibitor is currently tested in phase 2 clinical trials for non-small cell lung carcinomas (Gerber, 2019) and GBM (W. Kelly et al., 2020). In contrast to C75, TVB-2640 did not cause a decrease of basal invasion rate. Of note, the proliferation rate of LN-229 cells was not affected by the used concentrations of C75 and TVB-2640, indicating that the observed decreased invasiveness is not a result of decreased proliferation rate (Supplemetary Figure 4J). We thus decided to use the TVB-2640 inhibitor in further experiments in combination with formate. Efficacy of TVB-2640 on FASN was validated with ^13^C tracing, confirming that the compound effectively inhibits *de novo* fatty acid synthesis in our experimental set-up also in presence of formate (Figure 4H). We also confirmed that lower dose of TVB-2640 (1 μM) was already enough to prevent formate induced invasion (Supplementary Figure 4L-M).

To test if fatty acid synthesis is involved in transmitting the formate-dependent signaling towards increased MMP2 expression and activity in LN-229 cells, we analyzed MMP2 gene expression by RT-qPCR after TVB-2640 treatment (Figure 4I). In line with our hypothesis, we oberserved that cells treated with the FASN inhibitor TVB-2640 did not increase MMP2 expression upon formate.

To verify these findings in a more physiological model, we performed *ex vivo* brain slice assays and implanted LN-229 cells to test the formate-dependent effect on invasion after TVB-2640 treatment. As observed *in vitro*, TVB-2640 prevented the formate-induced increase in single cell velocity (Figure 4J). This suggests that pharmacologic inhibition of fatty acid synthesis can block the formate-induced pro-invasive effect in an *ex vivo* model.

In summary, these results indicate that formate promotes fatty acid synthesis to activate a pro-invasive cellular phenotype.

## DISCUSSION

Our results pinpoint to a physiological relevant role of formate to promote cancer cell invasion and metastasis. They substantiate earlier findings from our laboratory indicating that formate promotes cancer cell invasion *in vitro* (Meiser et al., 2018). While extracellular formate had no impact on migration of cancer cells, we found that it clearly has a promotive effect on invasion, which is in line with the notion that migration and invasion can be uncoupled (Schaeffer et al., 2014). Mechanistically, we uncovered that formate triggers fatty acid synthesis which promotes an invasive phenotype through proteolytic degradation of the extracellular matrix by MMPs. Furthermore, we found that formate does not only promote local invasion but does also increase the metastatic seeding capacity of breast cancer cells. Both, increased invasion and adhesion likely contributed to this observation.

In sum, this study broadens the current knowledge on the role of one-carbon metabolism and formate-overflow in the context of cancer progression. In that regard a question of particular interest is related to the physiological relevance of formate concentration. We could show here, that reduced invasion was observed upon suppression of formate release. This can be compensated by addition of 500 μM of exogenous formate. Previous findings of our lab showed that 100 μM of formate are already sufficient to promote cancer cell invasion (Meiser et al., 2018). While the plasma concentration of murine and human formate is in the range of 20–50 μM (https://hmdb.ca/metabolites/HMDB0304356; Meiser et al., 2018); first analyses of tumor interstitial fluid (TIF) from colorectal cancer (CRC) suggest that formate concentrations in the tumor microenvironment can increase significantly above 100 μM (Ternes et al., 2022). Based on these observations, we assume that the applied formate concentrations are likely to be within the physiological range of the tumor microenvironment. Further analysis to quantify the concentration of formate in different tumour tissues is currently ongoing.

Recent work in the context of CRC revealed that microbiome-derived formate and related high formate concentrations in the gut (10 mM) exacerbate CRC progression via AHR signaling (Ternes et al., 2022). Here, we provide evidence that significantly lower formate concentrations can already impact cancer cell invasion albeit following different mechanisms. Our transcriptome analysis in GBM cells strongly pinpoints towards a role of lipid metabolism. In conclusion, context-dependent and mechanistic understanding of the processes that underly formate-dependent activation of invasion are important to design effective intervention strategies in different tumor types.

To deepen our current understanding of how formate triggers downstream responses, more mechanistic insights are needed. Following our TF binding analysis (Supplementary Table 1), we identified some candidates, including the previously mentioned AHR (Ternes et al., 2022) and others related to immune-dependent or epigenetic roles. This list of identified candidates now serves for follow-up studies.

The here shown data also highlight that formate, in addition to its role as an anabolic substrate, can play a role as a signaling molecule in cellular plasticity. Such a fundamental function might also explain the observation that loss of *MTHFD1L* results in neural-tube defects during embryogenesis (Momb et al., 2013). Successfull formation of the neural tube requires proliferation, correct cell polarity and cell migration (Gilbert., 2000). While our data suggest a mechanism through which formate promotes invasion, it remains to be investigated whether formate signaling is essential for tissue organization in embryogenesis.

It has been demonstrated earlier that mitochondrial one-carbon metabolism is dispensable for cancer cell proliferation as the cytosolic part of one-carbon cycle can compensate for the loss of mitochondrial activity (Ducker and Rabinowitz, 2017). We can confirm these results, as we did not observe a growth reduction upon loss of *MTHFD1L*. Nevertheless, knockdown of *MTHFD1L* was shown to specifically inhibit formate overflow (Meiser et al., 2018; Meiser et al., 2016), which further argues for an additional, non-anabolic role of formate in cancer cell invasion. This hypothesis is substantiated by another recent study from our group in which we were able to demonstrate that loss of *MTHFD1L* expression in breast cancer cells resulted in decreased pulmonary metastasis formation in an syngeneic orthotopic breast tumor model (Kiweler et al., 2022). Interestingly, we and others found here that nucleotide synthesis inhibition by methotrexate (MTX) promotes formate overflow and that perturbation of mitochondrial serine catabolism reduces the migratory capacity of formate overflow positive, MTX-treated cells (Kiweler et al., 2022; Soflaee et al., 2022). However, in our current study, we observed that enhanced extracellular formate levels only enhance cell invasion, not cell migration. This suggests that the sustained migratory potential upon MTX depends on endogenous, mitochondrial serine catabolism and its associated biochemical outputs other than excreted formate. Hence, we hypothesize that the specific motility phenotype associated to mitochondrial serine catabolism can be impacted by two separate effects: (i) serine catabolism, and metabolic outputs thereof, and (ii) local concentrations of formate in the tumor microenvironment. In turn, such local formate concentrations depend on the rate of formate overflow and the fate of formyl-tetrahydrofolate within the mitochondrial one-carbon cycle. For example, increased expression of ALDH1L2 can lower formate overflow and vice versa. Similarly, oxidation via ALDH1L2 or hydrolysis via *MTHFD1L* either generates mitochondrial NADPH or ATP, respectively. This further illustrates the plasticity of serine catabolism and the different metabolic pathways that cancer cells can engage to adapt to changes within the local tumor microenvironment and along the metastatic cascade (Benzarti et al., 2020; Fendt et al., 2020).

In addition to the activation of MMPs, we uncovered that high extracellular formate levels promote fatty acid synthesis. Increased *de novo* fatty acid synthesis is an important determinant of tumor malignancy and presents thus an attractive target for clinical intervention (Menendez and Lupu, 2007). While *de novo* fatty acid synthesis is clearly associated with membrane biogenesis for cancer cell duplication, the mechanism by which this pathway regulates the processes of tumor cell invasion and migration is still not fully understood. GBMs occure in a lipid-rich environment as the dry weight of a human brain is composed to more than 50% of lipids and the brain itself contains about a quarter of the body’s cholesterol (O’Brien and Sampson, 1965). Since much of this lipid is housed in structural elements of neural tissue such as myelin, it is speculated that the amount of free lipids is relatively low which suggests that GBM cells must increase lipogenesis (Zhou and Wahl, 2019). The fatty acid synthesis genes ACC and FASN are both highly expressed in glioblastoma and have been associated with poor patient outcomes (Taïb et al., 2019). Furthermore, breast cancers that are capable of metastasizing to the brain showed evidence of altered lipid metabolism (Ferraro et al., 2021; Jin et al., 2020).

Based on this evidence, lipid metabolism has become an attractive target in studies of metabolic drivers of glioblastoma invasion and brain metastasis. Our data indicate that formate-dependent signaling could act as an upstream regulator of fatty acid synthesis to promote invasion and metastasis. Importantly, targeting fatty acid synthesis with the specific FASN inhibitor TVB-2640 was effective to inhibit formate-dependent invasiveness in *ex vivo* brain tissue slices. Notably, TVB-2640 is in clinical development for the treatment of CRC and very encouraging results with a 66.7% overall response rate have also been reported from a phase 1 trial of TVB-2640 in combination with bevacizumab in recurrent GBM (W. Kelly et al., 2020). Our results indicate fatty acid synthesis targeting as a mechanistic entry point to impede formate-dependent GBM invasion and to possibly also hinder metastatic progression of other tumor types. We have already demonstrated in different genetically engineered mouse models (GEMMs) that the extent of formate release rate depends on the oxidative state of the tumor (Meiser et al., 2018). It is thus tempting to speculate that especially in highly oxidative tumors with high formate overflow formate might act as an upstream regulator of fatty acid synthesis and that targeting FASN can be an effective strategy to block formate-dependent pro-invasive effects.

In summary, we pinpoint MMPs and FASN as two entry points to prevent formate induced cancer cell invasion. Further studies are however needed to increase insight into additional mechanisms that underly formate-induced cancer cell invasion and to explore how our findings can be exploited for therapeutic application. Certainly, such novel strategies to overcome cancer cell dissemination will be of utmost clinical importance.

## MATERIAL AND METHODS

### Chemicals

Sodium formate (**Sigma-Aldrich: 71539**), sodium acetate (**Sigma-Aldrich: S2889**), sodium lactate (**Sigma-Aldrich: 1614308**) and glycine (**Sigma-Aldrich: 50046**) were purchased from Sigma-Aldrich.

The MMP inhibitors: ARP100 (MMP2 inhibitor) (**Santa Cruz: SC-203522**) and MMP-9 Inhibitor I (MMP9 inhibitor) (**Santa Cruz: SC-311437**) were purchased from Santa Cruz. The FASN inhibitors, C75 (**Cayman Chemicals: 10005270-5**) and TVB-2640 (**Selleckchem: S9714**) were purchased from Cayman Chemicals and Selleckchem respectively.

### Tumor cells and culture conditions

The human glioblastoma LN-229 and U87 cells and the human breast cancer MDA-MB-468 cells were cultured in Dulbecco’s modified Eagle’s medium (DMEM) no glucose, no glutamine and no phenol red (**Thermo Fisher Scientific: A1443001**) supplemented with 17 mM glucose (**Sigma: G8769**), 2 mM glutamine (**Sigma-Aldrich: 25030081**) and 10% qualified south American fetal bovine serum (FBS) (**Thermo Fisher Scientific: 10500064**).

The human glioblastoma NCH601 cells (Schuster et al., 2020) were cultured as non-adherent spheres in DMEM-F12 medium (**Lonza: 12634010**) containing 1X Bit-100 (**Provitro: 2043100**), 2 mM of glutamine (**Sigma-Aldrich: 25030081**), 30 U/ml Pen/Strep (**Westburg: LO DE17-602E**), 20 ng/ml BFGF (**Miltenyi: 130-093-841**), 1 U/ml heparin (**Sigma-Aldrich: H0200000**) and 20 ng/ml EGF (**Provitro: 1325950500**). shScramble and sh*MTHFD1L* have been described in (Meiser et al., 2018).

The human glioblastoma BG5, BG7, GG6 and GG16 (Schuster et al., 2020) were cultured as non-adherent spheres in Neurobasal base medium (**Thermo Fisher Scientific: 21103049**) containing 1x B27 without vitamine A (**Thermo Fisher Scientific: 12587010**), 2 mM glutamine (**Sigma-Aldrich: 25030081**), 30 U/ml Pen/Strep (**Westburg: LO DE17-602E**); 20 ng/ml BFGF (**Miltenyi: 130-093-841**), and 20 ng/ml EGF (**Provitro: 1325950500**).

To reflect human physiological conditions, the cells were cultured in Plasmax medium. Plasmax medium was self-made following the protocol in Vande Voorde *et al*. (Vande Voorde et al., 2019). Chemicals used to prepare the medium are listed below:

| Chemical Name | Concentration (μM) |  |
| --- | --- | --- |
| **Proteinogenic Amino Acids** |  |  |
| L-Alanine | 510 | Carl Roth: 3076.1 |
| L-Arginine monohydrochloride | 65 | Carl Roth : 1689.1 |
| L-Asparagine | 41 | Sigma-Aldrich: A0884 |
| L-Aspartate | 6 | Sigma-Aldrich: A9256 |
| L-Cysteine hydrochloride | 33 | Carl Roth: 3468.1 |
| L-Glutamate | 98 | Sigma-Aldrich: G1251 |
| L-Glutamine | 650 | Thermo Fisher Scientific: 25030081 |
| Glycine | 330 | Sigma-Aldrich: 50046 |
| L-Histidine monohydrochloride monohydrate | 120 | Applichem: A3733 |
| L-Isoleucine | 140 | Carl Roth : 1698.1 |
| L-Leucine | 170 | Carl Roth : 1699.1 |
| L-Lysine | 220 | Sigma-Aldrich: L5626 |
| L-Methionine | 30 | Carl Roth : 1702.1 |
| L-Phenylalanine | 68 | Carl Roth : 1709.1 |
| L-Proline | 360 | Sigma-Aldrich: 81709 |
| L-Serine | 140 | Sigma-Aldrich: 84959 |
| L-Threonine | 240 | Carl Roth : 1738.1 |
| L-Tryptophan | 78 | Carl Roth : 1739.1 |
| L-Tyrosine | 74 | Sigma-Aldrich: T3754 |
| L-Valine | 230 | Carl Roth : 1742.1 |
| **Non-proteinogenic Amino Acids** |  |  |
| L-2-Aminobutyric acid | 41 | Sigma-Aldrich: A2536 |
| L-Citrulline | 55 | Sigma-Aldrich: C7629 |
| L-Cystine | 65 | Sigma-Aldrich: C7602 |
| L-Homocysteine | 9 | Sigma-Aldrich: 69453 |
| Trans-4-Hydroxy-L-proline | 13 | Sigma-Aldrich: 56250 |
| L-Ornithine monohydrochloride | 80 | Carl Roth: 1703.1 |
| L-Pyroglutamate | 20 | Sigma-Aldrich: 83160 |
| **Amino Acids Derivates** |  |  |
| N-Acetyl glycine | 70 | Sigma-Aldrich: A16300 |
| L-Carnosine | 6 | Carl Roth : 4148.2 |
| L-Glutathione (reduced) | 37 | Carl Roth : 6832.2 |
| Taurine | 130 | Sigma-Aldrich: T0625 |
| Betaine | 72 | Sigma-Aldrich: 61962 |
| **Other Components** |  |  |
| Sodium Acetate | 42 | Sigma-Aldrich: S2889 |
| Acetone | 55 | Sigma-Aldrich: 34850 |
| Acetyl carnitine | 5 | Sigma-Aldrich: A6706 |
| Citrate | 114 | Sigma-Aldrich: 27488 |
| L-Carnitine hydrochloride | 46 | Sigma-Aldrich: C0283 |
| Creatine | 37 | Sigma-Aldrich: C0780 |
| Creatinine | 74 | Sigma-Aldrich: C4255 |
| Formate | 33 | Sigma-Aldrich: 71539 |
| D-Glucose | 5560 | Sigma-Aldrich: G8769 |
| Glycerol | 82 | Sigma-Aldrich: G2025 |
| 2-Hydroxybutyric acid sodium salt | 31 | Sigma-Aldrich: 220116 |
| (R)-(-)-3-Hydroxybutyric acid sodium salt | 77 | Sigma-Aldrich: 298360 |
| (±)-Sodium β-hydroxyisobutyrate | 20 | Sigma-Aldrich: 36105 |
| Hypoxanthine | 5 | Sigma-Aldrich: H9377 |
| Sodium L-Lactate | 500 | Sigma-Aldrich: 71718 |
| Methyl acetoacetate | 41 | Sigma-Aldrich: 537365 |
| Phenol Red sodium salt | 25 | Carl Roth : 2566.1 |
| Sodium pyruvate | 100 | Sigma-Aldrich: S8636 |
| Succinic acid sodium salt | 23 | Sigma-Aldrich: 224731 |
| Uracil | 2 | Sigma-Aldrich: U0750 |
| Uric acid | 270 | Carl Roth : 4999.1 |
| Urea | 3000 | Carl Roth : 7638.1 |
| Uridine | 3 | Carl Roth : 0714.1 |
| **Inorganic Salts** |  |  |
| Ammonium Chloride | 50 | Sigma-Aldrich: A9434 |
| Calcium Chloride | 1800 | Carl Roth : CN93.1 |
| Magnesium Sulfate | 813 | Carl Roth : 0682.1 |
| Potassium Chloride | 5330 | Sigma-Aldrich: P9541 |
| Sodium Bicarbonate | 26191 | Sigma-Aldrich: S6297 |
| Sodium Chloride | 118706 | Carl Roth : HN00.2 |
| di-Sodium hydrogen phosphate heptahydrate | 1010 | Carl Roth : X987.1 |
| **Trace Elements** |  |  |
| Ammonium monovanadate | 0.0026 | Carl Roth : 4232.1 |
| Copper(II) sulphate | 0.0052 | Carl Roth : CP86.1 |
| Iron(III)nitrate nonahydrate | 0.1238 | Sigma-Aldrich: 216828 |
| Iron(II) sulphate heptahydrate | 1.0428 | Carl Roth : P015.1 |
| Manganese(II) chloride tetrahydrate | 0.0002 | Carl Roth : T881.3 |
| Sodium Selenite | 0.0289 | Sigma-Aldrich: S5261 |
| Zinc sulphate heptahydrate | 1.5 | Carl Roth : K301.1 |
| **Vitamins** |  |  |
| Ascorbic acid | 62 | Sigma-Aldrich: A0278 |
| BME Vitamins Solution 100x | 1x | Sigma-Aldrich: B6891 |
| Vitamine B12 | 0.005 | Sigma-Aldrich: V6629 |

The cultures were maintained at 37°C in a humidified 5% CO_2_ /95% air atmosphere. Cell lines were regularly tested for mycoplasma contamination. No contaminations have been encountered in the period during which the experiments were carried out.

LN-229 cells and MDA-MB-468 cells were authenticated by eurofins (ID:11107531009).

### Transwell Migration/Invasion Assay

The *in vitro* migration potential was determined on non-coated transwell chambers with 8 μm pore size (**Greiner Bio-one: 662638**). To assess the cell’s ability to invade, the same transwell chambers were coated with a collagen-Extracellular matrix (ECM) coating (standard coating). Compounds to prepare the coatings are listed below:

| Coating | Composition |
| --- | --- |
| Collagen-ECM | 0.05 mg/ml collagen type I (**Sigma-Aldrich: A1048301**), 0.5 mg/ml protein of ECM gel (**Sigma-Aldrich: E1270**) |
| Collagen | 0.05 mg/ml collagen type I (**Sigma-Aldrich: A1048301**) |
| ECM | 0.5 mg/ml protein of ECM gel (**Sigma-Aldrich: E1270**) |
| Fibronectin | 1.12 µg/ml protein of ECM gel (**Sigma-Aldrich: F0895**) |
| Laminin | 0.01 mg/ml Laminin from murine sarcoma basement membrane (**Sigma-Aldrich: L2020**) |
| Poly-L-lysine (PLL) | 25 µg/ml Poly-L-lysine hydrobromide (**Sigma-Aldrich: P1274**) |

In brief, 300 μl of cell solution without FBS (containing 5 x 10^4^ cells) were pipetted into the transwell chamber in duplicates per condition. 750 μl of DMEM supplemented with 10 % FBS, which acts as chemoattractant, was added into the bottom of the lower chamber. In case of chemical interventions, the compound was spiked into the upper chamber. After 24 h, the cells were fixed with 4% of Formaldehyde (**VWR: 11699404**) and stained with 0.05% of crystal violet solution (**Sigma-Aldrich: HT90132**) for 15 minutes, respectively. Non-invading cells were removed at the top of the chamber with a cotton tip and invasion was quantified by counting the number of cells per chamber. Five pictures/chamber were taken using an inverted microscope at a 20x magnification. The cell count was assessed using a Python script written by the AI department of Helmholtz Muenchen. The Python code and full description of the tool can be accessed in github (https://github.com/HelmholtzAI-Consultants-Munich/Automatic-Cell-Counter) (Piraud, 2022).

### *Ex vivo* Brain Slice Invasion Assays

The *ex vivo* brain slice assays were performed as in Schuster *et al*., (Schuster et al., 2020). In brief, NSG mice (NOD.Cg-Prkdc^SCID^ Il2rg^tm1Wjl^/SzJ) were sacrificed by cervical dislocation, and the brains were rapidly transferred to an ice-cold cutting solution (0.1% of GlutaMax (**Thermo Fisher Scientific: 35050061**), 25 mM of HEPES (**Thermo Fisher Scientific: 15630080**), 50 U/ml Pen-Strep (**Westburg: LO DE17-602E)** in DMEM). Employing the McIIwain tissue chopper, 400 μm thick brain slices were generated. Up to two slices were cultured on a 1 μm transwell chamber (**Greiner Bio-one: 657631**) in 1 ml Hibernate™-A medium (**ThermoFisher: A1247501**) supplemented with 20% BIT-100 (**Provitro: 2043100**) and 100 U/ml Pen-Strep (**Westburg: LO DE17-602E**).

The brain slices were cultivated at 37°C and 5% CO_2_ up to 3 days.

Subsequently, 5 x 10^4^ GFP-labeled tumor cells, contained in a maximal volume of 1 μl of ECM, were manually injected above the corpus callosum. The culture medium containing the treatment was exchanged every second day. After 10 days, live cell imaging using the IncuCyte® Live-Cell Analysis system was performed over 24 h (**Essen Bioscience**) (Green channel, x4 objective, a picture was taken every 2 h) and the velocity of the single cells was assessed with ImageJ (15 cells per condition and 8 replicates per cell line).

### *In vivo* Work

#### Orthotopic brain tumor model

To assess the invasion ability of the NCH601 cells *in vivo*, the cells (sh*Scramble* or *MTHFD1L* KD) were injected intracranially into the right frontal part of the brain according to the protocol published in Oudin A. *et al., (Oudin et al., 2021)*. Briefly, NU/NU Nude female mice (n=16) were anesthetized and 3 x 10^5^ NCH601 cells were implanted into the right hemisphere of the brain. Two or four months after tumor cell implantation, animals were perfused with cold heparinized saline solution followed by fixative solution (formol 4% buffered, formaline 10%) under deep anesthesia. The brains were harvested, placed in sucrose gradient solution (30%) for 48h and stored at −80°C. Brains were cryosectioned coronally (10 μm), mounted on slides, and stained with 4′,6-diamidino-2-phenylindole (DAPI) (**ThermoFisher Scientific: D1306**) according to the manufacturer’s protocol. Intracranial distribution of GFP-positive-NCH601 cells was assessed with fluorescent microscopy (x4 objective). Invasion was finally obtained by: LogRatio(Fluorescent Signal Left Hemisphere/Fluorescent Signal Right Hemisphere). Quantification was assessed using a Fiji (DOI: 10.1038/nmeth.2019) plugin written by Dr. Aymeric FOULQUIER D’HEROUEL from the University of Luxembourg (Fouquier d’Hérouël, 2021). The code and full description of the tool can be accessed in github (https://github.com/monomeric/BrainSignalDistribution).

#### Tail vein injections

To assess the seeding capacity of the MDA-MB-468 cells *in vivo*, 1 million cells (sh*Scramble*, 24 h formate pre-treated cells or *MTHFD1L* KO) were injected intravenously in the NSG mouse (NOD.Cg-Prkdc^SCID^ Il2rg^tm1Wjl^/SzJ) tail vein. The experiment was terminated after six weeks and lung und liver were prepared for examination of metastatic outgrowth. No metastases were found in the lung. Macroscopic liver metastases were blindly counted under a microscope and microscopic liver metastases were visualized by H&E staining.

#### H&E staining

For H&E staining, snap-frozen livers were embedded in Tissue-Tek® O.C.T (**Sakura: 4583**) and cut in 10 μm sections using Leica Cryostat (**Leica Biosystems**). 5 sections in a distance of 100 μm to each other were stained per liver. Staining was performed according to established protocols. Briefly, selected sections were dehydrated with MeOH (**Carl Roth: 83885)** and stained with Gill 2 hematoxylin (**Sigma-Aldrich: GHS232)**. Sections were neutralized with successive washes of tap water, hard water (10g MgSO4 (**Carl Roth: 0682.1)** and 0.7g NaHCO_3_ (**Sigma-Aldrich: S6297)** per L), and distilled water. Afterwards, sections were stained with Eosin-solution (**Sigma-Aldrich: 1.09844)** and dehydrated through successive washes with 80%, 95%, 100% EtOH. Before mounting on a glass slide, the liver were stained with Xylol (**VWR: 2875.325**).

#### RNA-Seq

Total RNA was extracted directly from the cell culture dishes using RNeasy Mini Kit (**Qiagen: 74104**). RNA integrity and quantity was determined using the Agilent 2100 Bioanalyzer according to the manufacturer’s manual. A RNA Integrity Number (RIN) above 9 was accepted for downstream analysis.

For library preparation, 1 μg of RNA was poly-(A) selected, fragmented, and reverse transcribed by using the Elute, Prime, and Fragment Mix from Illumina.

End repair, A-tailing as well as adaptor ligation and library enrichment were performed as described in the TruSeq Stranded mRNA library preparation kit (**Illumina: RS-122-2101**) according to the manufacturers manual. RNA libraries were finally assessed for quality with the Agilent 2100 BioAnalyzer using DNA high sensitivity kit (**Agilent: 5067-4626**) and quantified using Qubit DNA high sensitivity assay (**Thermo Fischer Scientific: Q32854**). Eqimolar pool of all RNA libraries were sequenced as 75 bp single-read run on an Illumina NextSeq500 sequencer at LCSB sequenicng platform (**RRID: SCR_021931**).

RNA-Seq Raw Data was uploaded on GEO under the submission number:GSE196706.

#### RNA Extraction, cDNA Synthesis, and qPCR Analysis

Total RNA was extracted directly from the cell culture dishes using the RNeasy Mini Kit (**Qiagen: 74104**). Then, the extracted RNA was converted to cDNA using the High-capacity cDNA Reverse Transcription Kit (**Thermo Fisher Scientific: 4368814**). RT-qPCR was performed from 20 ng of cDNA per sample using Fast SYBR™ Green Master Mix (**Thermo Fisher Scientific: 4385614**). All samples were analyzed in technical triplicates. qPCR using Fast SYBR Green was conducted at 95°C for 20 seconds, followed by 40 cycles of 95°C for 1 second and finally 60°C for 20 seconds. The comparative Ct method was used to quantify the relative mRNA levels. Experiments were performed using the QuantStudio 5 Real-Time PCR System (**Applied Biosciences, ThermoFisher Scientific**) and data processing was performed using QuantStudio Design&Analysis v1.5.1 software (**Applied Biosciences, ThermoFisher Scientific**). All expression data was normalized to two housekeeping controls (*GAPDH* and *CycloA*). The primers used in this study are listed in the table below:

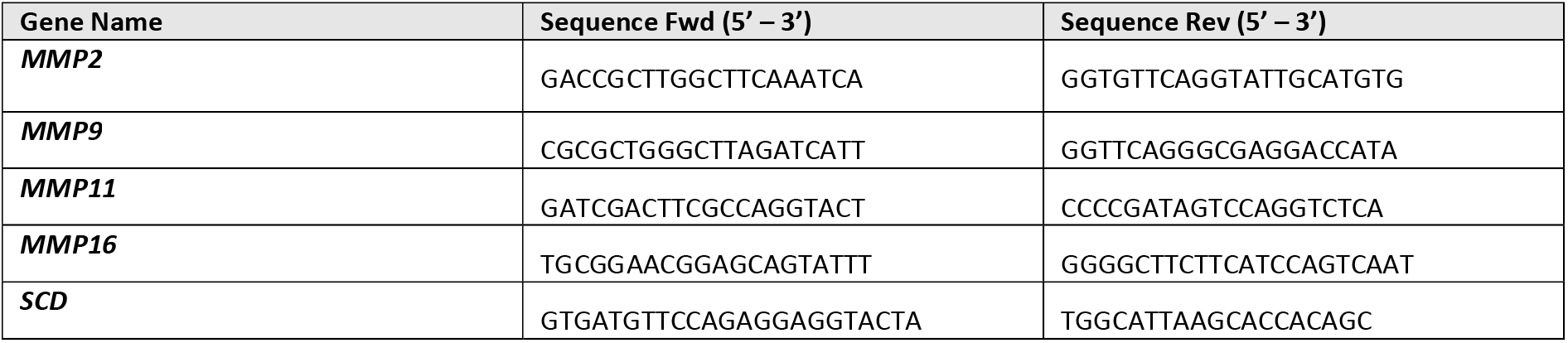
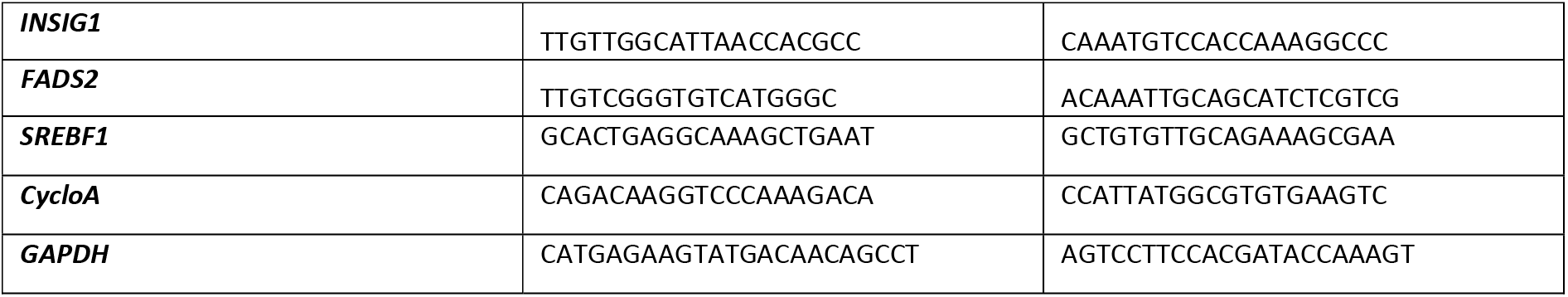

#### Western Blot

Total cell lysates were extracted from adherent cells by incubating the cells for 10 minutes on plate in ice cold cell lysis buffer (150 mM NaCl (**Sigma-Aldrich: S3014**); 1 mM EDTA (**Sigma-Aldrich: ED**); 50 mM Tris-HCl (**Carl Roth: 0188.2**); 1% IGEPAL^®^ CA-630 (**Sigma-Aldrich: I3021**) containing Protease and Phosphatase Inhibitors (**Sigma-Aldrich: 490687001 & 4693124001)**). Lysates were sonicated for 10 minutes to increase cell lysis efficiency. Purification of the lysate was achieved by centrifugation (13,000g, 10 minutes at 4°C).

To asses MMP abundance in the cell culture medium, the cells were cultured for 72 h on a collagen-ECM-coating ((0.05 mg/ml collagen type I (**Sigma-Aldrich: A1048301**), 0.5 mg/ml protein of ECM gel (**Sigma-Aldrich: E1270**)) in medium without FBS. Proteins contained in the medium were concentrated using Pierce™ Protein Concentrator PES tubes (**Thermo Fisher Scientific: 88517**).

Protein concentration was assessed by Bradford assay. 30 μg of total protein were loaded on RunBlue 4-12% Bis-Tris gels (**GeneScript: M00656**) supplemented with 4x NuPage LDS Sample buffer (**Thermo Fisher Scientific: NP0007**) and 10 mM DTT (**Sigma-Aldrich: 10708984001**). After migration, proteins were blotted on nitrocellulose membrane according to standard protocols.

Total protein loading was quantified using REVERT staining solution (**LI-COR: 926-11011**) and imaged with the Odyssey CLx Imaging System (**LI-COR**) at 700 nm. Blocking of the membrane was carried in 5% milk solution for 1 hour at room temperature. Primary antibodies were incubated at 4°C over night at a 1:1000 dilution, followed by incubation with the secondary antibody for 1 hour at room temperature. Detection of the fluorescent signal was performed with the Odyssey CLx Imaging System (**LI-COR**). Image analysis was done using the ImageStudioLite Software Vers.5.2 (**LI-COR**).

| Antibody | Reference |
| --- | --- |
| β-actin | Cell Signaling Technology (3700) clone 8H10D10, lot18 |
| AMPKα (D5A2) | Cell Signaling Technology (5831) clone D5A2, lot8 |
| Phospho-AMPKα (Thr172) (40H9) | Cell Signaling Technology (2535) clone T172, lot21 |
| MMP-2 (D4M2N) | Cell Signaling Technology (40994) clone D4M2N, lot3 |
| MMP-9 | Abcam (ab38898) clone GR3204084, lot24 |
| Acetyl-CoA Carboxylase | Cell Signaling Technology (3676) clone C83B10, lot9 |
| Phospho-Acetyl-CoA Carboxylase (Ser79) (D7D11) | Cell Signaling Technology (11818) clone D7D11, lot6 |
| mTOR (7C10) | Cell Signaling Technology (2983S) clone 7C10, lot14 |
| Phospho-mTOR (Ser2448)(D9C2) | Cell Signaling Technology (5536S) clone D9C2, lot5 |

#### Gelatin Zymography

To asses matrix metalloproteinase activity, the cells were cultured for 72 h on a collagen-ECM-coating ((0.05 mg/ml collagen type I (**Sigma-Aldrich: A1048301**), 0.5 mg/ml protein of ECM gel (**Sigma-Aldrich: E1270**)) in medium without FBS.

Subsequently, the proteins in the medium were concentrated using Pierce™ Protein Concentrator PES tubes (**Thermo Fisher Scientific: 88517**). Protein concentration was determined by Bradford assay. 5 μg of total protein were loaded on NovexTM 10% Zymogram Plus (Gelatin) Protein Gels (**Thermo Fisher Scientific: ZY00102BOX**) using 2x non-reducing sample buffer (1M Tris-HCl pH 6,8 (**Carl Roth: 0188.2**); 10% SDS (**Carl Roth: 0183.3**); 50% glycerol (**Sigma-Aldrich: G5516**), 0.1% bromophenol blue (**Bio-Rad: 1610404**)). After electrophoresis the gels were washed in a 2.5% Triton X100 solution (**Carl Roth: 3051.3**) and incubated overnight in an incubation buffer (1% Triton X-100 (**Carl Roth: 3051.3**); 50 mM Tris-HCl pH 7.5 (**Sigma-Aldrich: C1016**); 1 μM Zinc sulphate heptahydrate (**Carl Roth: K301.1**)) at 37°C for the development of zymolytic bands. Protease bands were detected by absence of Coomassie Brilliant Blue staining as a result of digested gelatin. The bands were quantified using ImageJ.

#### Adhesion Assay

Coverslips were ECM-collagen-coated prior to seeding 50,000 cells per cover slip. The cells (Ctrl. *vs* 24h formate pre-treated cells) could adhere to the cover slips for 1 hour prior fixation. The cells were fixed with 4% Formaldehyde (**VWR: 11699404**) and stained with 0.05% crystal violet solution (**Sigma-Aldrich: HT90132**) for 15 minutes, respectively. Five pictures/cover slip were taken using an inverted microscope at a x20 magnification. The cell count was assessed using a Python script written by the AI department of Helmholtz Muenchen. The Python code and full description of the tool can be accessed in github (https://github.com/HelmholtzAI-Consultants-Munich/Automatic-Cell-Counter) (Piraud, 2022).

#### Stable Isotope tracing and metabolite extraction

Stable isotope tracing experiments with [U-^13^C]-glucose tracer (**Cambridge Isotope Laboratories: CLM-1396**) were performed in DMEM without glucose, glutamine and phenol red, supplemented with 2 mM glutamine, 17 mM glucose tracer, and 10 % FBS or 10% charcoal stripped FBS (**Sigma-Aldrich: F6765**).

1.5 x 10^5^ LN-229 cells were seeded in triplicates in 12-well plates. 24 h prior the experiment, the cells were cultivated in medium without [U-^13^C]-glucose tracer. To normalize, identical triplicate wells were seeded and cell size and cell count was assessed in parallel to metabolite extraction. Packed cell volume (PCV) at the end of the experiment was used to normalize intracellular metabolite levels.

After culturing the cells for 24 h (or 72 h for fatty acid detection) in the tracer medium, metabolites were extracted for GC-MS analysis. To do so, the cells were washed with ice cold 0.9% NaCl solution (**Carl Roth: HN00.2**). 400 μl ice-cold MeOH (**Carl Roth: KK44.1**)/H2OMQ ((ratio, 1:1) containing the internal standards pentanedioc-d6 acid and [U-^13^C]-ribitol at a final concentration of 1 μg/ml and Tridecanoid-d25 acid at a final concentration of 5 μg/ml) were added to each well. The plates were then incubated for 5 minutes at 4°C on a rocking shaker. Supernatant was collected and transferred into Eppendorf tubes containing 200 μl ice-cold chloroform (**Sigma-Aldrich: 154733**). After centrifugation at 13,000 g, for 10 minutes at 4°C, the sample become biphasic.

Polar phase (or the unpolar phase for fatty acid detection) were transferred into GC-MS vials and dried in a speedvac at 4°C prior MS analysis.

To extract for LC-MS analysis, we cultured the cells for 24 h in the tracer medium. To do so, the cells were washed with ice cold PBS solution. LC-MS analysis was performed as described in (Meiser et al., 2016)

#### GC-MS analysis: Polar Phase

As described in (Kiweler et al., 2021). In brief, polar metabolites were derivatized for 90 min at 55 °C with 20 μl of methoxyamine (c = 20 mg/ml) in pyridine under continuous shaking and subsequently for 60 min at 55 °C with 20 μl of MTBSTFA w/1% TBDMCS. GC-MS analysis was performed using an Agilent 7890B GC coupled to an Agilent 5977A Mass Selective Detector (**Agilent Technologies**). A sample volume of 1 μl was injected into a Split/Splitless inlet, operating in splitless mode at 270 °C. Gas chromatograph was equipped with a 30 m (I.D. 250 μm, film 0.25 μm) ZB-35MS capillary column with 5 m guard column (**Phenomenex**). Helium was used as carrier gas with a constant flow rate of 1.2 ml/min. GC oven temperature was held at 100 °C for 2 min and increased to 300 °C at 10 °C/min and held for 4 min. Total run time was 26 min. Transfer line temperature was set to 280 °C. Mass selective detector (MSD) was operating under electron ionization at 70 eV. MS source was held at 230 °C and the quadrupole at 150 °C. For precise quantification of the MID, measurements were performed in selected ion monitoring mode.

The MetaboliteDetector software package (Version 3.220180913) was used for mass spectrometric data post processing, quantification, MID calculations, correction of natural isotope abundance, and determinations of fractional carbon contributions (DOI:10.1021/ac802689c).

#### GC-MS analysis: Non-Polar Phase

Derivatization was performed by using a multi-purpose sample preparation robot (GERSTEL). Dried non-polar extracts were dissolved in 30 μl N-tert-Butyldimethylsilyl-N-methyltrifluoroacetamide with 1% tert-Butyldimethylchlorosilane (MTBSTFA w/1% TBDMCS) and incubated for 60 min at 55 °C under continuous shaking.

GC-MS analysis was performed by using an Agilent 7890A GC coupled to an Agilent Mass Selective Detector (**Agilent Technologies**). A sample volume of 1 μl was injected into a Split/Splitless inlet, operating in splitless mode at 280 °C. The gas chromatograph was equipped with a 30 m (I.D. 0.25 mm, film 0.25 μm) ZB-5MSplus capillary column (**Phenomenex**) with 5 m guard column in front of the analytical column. Helium was used as carrier gas with a constant flow rate of 1.4 ml/min. GC temperature program: 100 °C for 1 minute then increased to 325 °C at 7.5 °C/min and held for 4 min. The total run time was 35 min. The transfer line temperature was set to 280 °C. The MSD was operating under electron ionization at 70 eV. The MS source was held at 230 °C and the quadrupole at 150 °C. GC-MS measurements were performed in selected ion monitoring mode for precise determination of the mass isotopomer distributions (MIDs).

#### Data processing and normalization

All GC-MS chromatograms were processed using MetaboliteDetector, v3.2.20190704 (Hiller et al., 2009). Compounds were annotated by retention time and mass spectrum using an in-house mass spectral (SIM) library (overall similarity: >0.85). The following deconvolution settings were applied: Peak threshold: 2; Minimum peak height: 2; Bins per scan: 10; Deconvolution width: 8 scans; No baseline adjustment; Minimum 1 peaks per spectrum; No minimum required base peak intensity. MIDs for the following fragments were calculated:

| Derivative | Metabolite | m/z - Range | Sum Formula (Fragment) | MID size |
| --- | --- | --- | --- | --- |
| Hexadecanoic acid 1TBDMS | C16:0 | 313-332 | C18H37O2Si | 17 |
| Hexadecenoic acid 1TBDMS | C16:1 | 311-330 | C18H35O2Si | 17 |
| Cholesterol 1TBDMS | Cholesterol | 443-473 | C29H51OSi | 28 |

#### LC-MS analysis

The LC-MS setup consisted in a Vanquish Flex (**Thermo Fisher Scientific**) liquid chromatography configured in binary gradient and coupled with a Q-Exactive Plus mass spectrometer (**Thermo Fisher Scientific**). The analytical column was a SeQuant ZIC-pHILIC (**Merck**) (2.1 mm × 150 mm, 5 μm) with a guard column (2.1 mm × 20 mm) and was heated at 45°C in the column oven. The solvent A consisted of 20 mM ammonium carbonate at pH 9.2 with 5 μM of medronic acid. The solvent B consisted in pure acetonitrile. Samples were injected onto the column and eluted by linear gradient ranging from 80% B to 20% B in 15 min at a constant flow rate of 200 μl/min. The MS acquisition was performed with a polarity switch between the positive and negative electrospray ionization modes. Full MS spectra were acquired from 75 to 1000 m/z at 70,000 resolution (at 200 m/z) with an automatic gain control set to 1e6 charges and a maximum ion trap fill was set to 250 ms.

#### Cell proliferation

Cell proliferation was determined using the IncuCyte^®^ Live-Cell Analysis system (**Essen Bioscience**). 5 x 10^3^ cells/well were seeded in 96-well plates and imaged every two hours (Brightfield, x10 objective, during 72h). Cell proliferation were determined as measurement of cell density (confluence) using the Incucyte^®^ analysis software.

#### Scratch Assay

Cell migration was determined using the IncuCyte® Scratch Wound Assay system for 96 well plates (**Essen Bioscience**). In brief, 5 x 10^4^ cells/well were seeded on a PLL coated (25 μg/ml Poly-L-lysine hydrobromide (**Sigma-Aldrich: P1274**)) 96 well plate. A wound was scratched in the cell monolayer across each well with the 96-pin IncuCyte WoundMaker Tool (**Essen Bioscience**) and the different treatments were applied to the cells. Measurement for individual treatment conditions was performed in biological replicates (n = 6). Confluence of the wound area was monitored every two hours for a total of 72 h. Cell migration was determined as measurement of cell density (confluence) in the wound area relative to the outer spatial cell density using the Incucyte^®^ analysis software.

#### Oxygen Consumption Rate

Basal Oxygen Consumption Rate (OCR) was measured using the XF96 Extracellular Flux Analyzer (**Agilent**) according to the manufacturer’s manual. Briefly, 4 x 10^3^ cells were seeded on a PLL coated (25 μg/ml Poly-L-lysine hydrobromide (**Sigma-Aldrich: P1274**)) Seahorse plate. Formate treatment was performed 24 h before the Seahorse measurement. The cells were sequentially treated with 1.5 μM oligomycine (**Sigma-Aldrich: 495455**); 15 μM of Carbonyl cyanide *m*-chlorophenyl hydrazone (CCCP)(**Sigma-Aldrich: C2759**); and a mixture containing rotenone (**Sigma-Aldrich: R8875**) and antimycin A (**Sigma-Aldrich: A8674**). Seahorse XFe Wave Software (**Agilent**) was applied to analyze the data. Protein concentration, necessary for normalization, was obtained by Bradford assay.

#### Statistics

Unpaired *t-*test with Welch’s correction was applied for pairwise comparison (two-sided) using GraphPad Software Vers.9. For normalization, data points of one experiment were either normalized to the untreated control or divided by the global mean of the individual experiment. We define one *n* as one independent experiment (in some cases further consisting of several wells, e.g. triplicate wells for all stable isotope tracing experiments). The mean of one independent experiment was considered as one *n*. The mean values of several independent experiments (as indicated in figure legends) were plotted and used for statistical analysis as indicated.

## Data availability

The data that support the findings of this study are available from the corresponding author upon reasonable request.

RNA-Seq Raw Data was uploaded on GEO under the submission Number:GSE196706

## Resource availability

Further information and requests for resources and reagents should be directed to and will be fulfilled by the Lead Contact, Johannes Meiser

## Author Contributions

Conceptualization, J.M., C.D.; Methodology, J.M., C.D., N.K., V.I.P., A.O., A.S.; C.J., S.P.N., E.L.; Software, A.F.H., R.S., M.P., A.S.; Validation, C.D., N.K., L.N., J.M.; Formal Analysis, C.D., N.K., V.I.P., L.N., C.J.; Investigation, C.D., N.K., V.I.P, L.N., A.O., V.B. E.L.; Resources, R.H., A.F.H., R.S., N.I.L., C.O., M.R., C.J., S.P.N., J.M., E.L.; Data Curation, R.H.; Writing – Original Draft, C.D., J.M.; Writing – Review & Editing, All Authors; Visualization, C.D., J.M., Supervision, J.M., M.P., A.S., E.L., S.P.N.; Project Administration, J.M.; Funding Acquisition, J.M., N.K., M.P., A.S., E.L., S.P.N.

## Acknowledgment

We thank Rolf Bjerkvig for providing BG5, BG7, GG6, GG16 GBM cells, Christel Herold-Mende for providing the NCH601 GBM cell line and Clément Thomas (LIH, Luxembourg) for providing 4T1 cells. We would also like to thank Virginie Baus (LIH, Luxembourg) for cutting the mouse brains. We would like to thank: the LCSB Metabolomics Platform, especially Xiangyi Dong and Floriane Vanhalle, for providing technical and analytical support. Also Francois Bernardin for technical asistance with Mass Spec analysis at LIH. We also thank all our collaboration partners for fruitful discussions and constructive feedback. All graphical figures were produced with BioRender.com. J.M. is supported by the FNR-ATTRACT program (A18/BM/11809970) and CORE program (C21/BM/15718879). N.K. is supported by the LIH Career Launchpad program (Legs Baertz) and by a DFG fellowship (KI 2508/1-1). AS and SPN are grateful for the financial support by Fondation Cancer Luxembourg (INVGBM project).

---

**Supplementary Figure 1.**
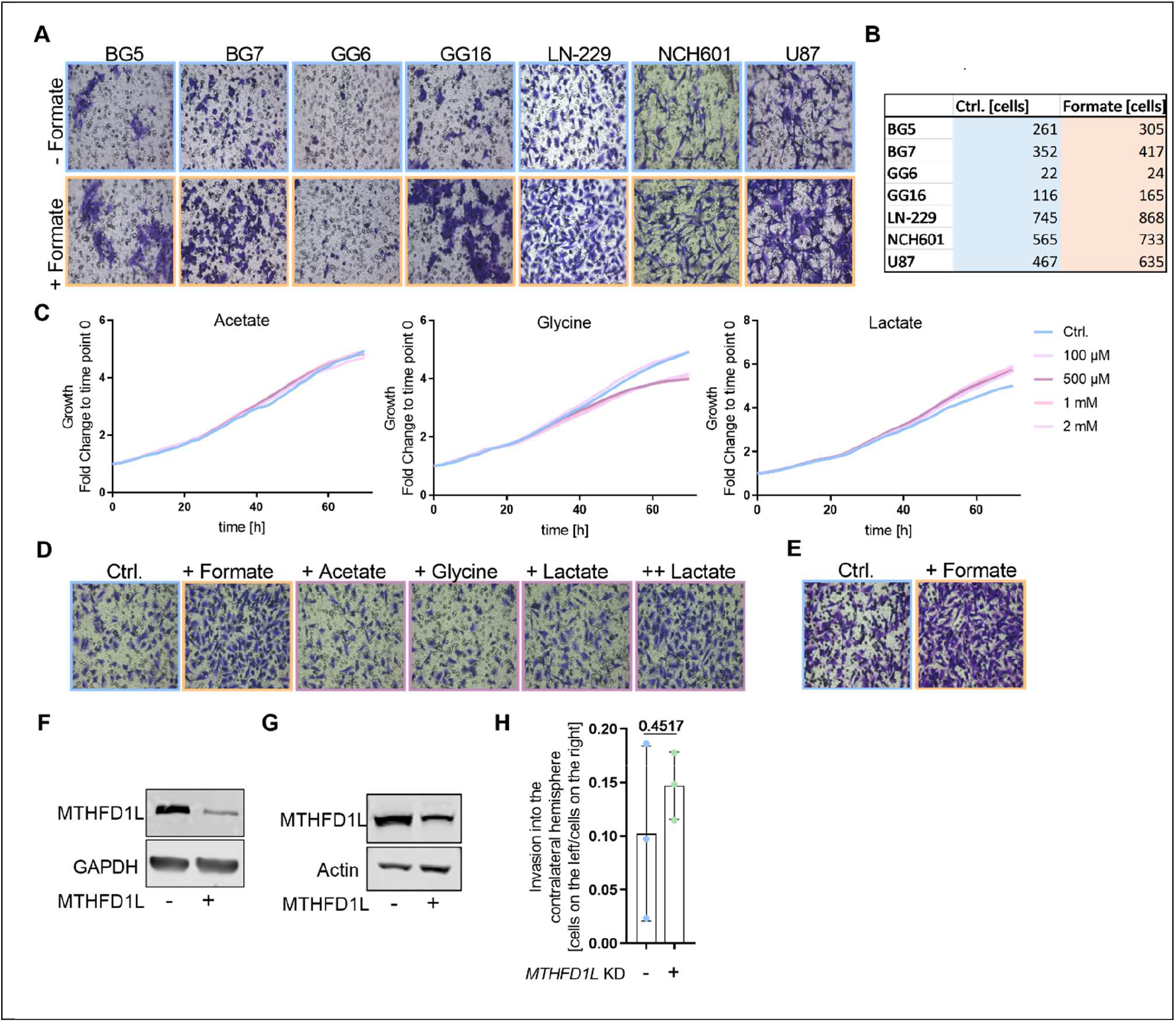
**(A)** Representative pictures of the Boyden chamber assay of different glioblastoma cells (BG5, BG7, GG6, GG16, LN-229, NCH601, and U87). **(B)** Average cell number per chamber for different glioblastoma cell lines. **(C)** Proliferation curves of LN-229 cells that were treated for 72 hours with different concentrations (0 μM, 100 μM, 500 μM, 1mM, 2mM) of small molecules (Na-formate, Na-acetate, glycine and Na-lactate). Representative curve is displayed as a fold change to time point zero. **(D)** Representative pictures of Boyden chamber assay of LN-229 cells which were treated with Na-formate, Na-acetate, glycine, and Na-lactate. **(E)** Representative pictures of Boyden chamber assay of LN-229 cells which were cultured in Plasmax medium. **(F)** Representative Western Blot showing MTHFD1L expression in LN-229 control and *MTHFD1L* KD cells. **(G)** Representative Western Blot showing MTHFD1L expression in NCH601 control and *MTHFD1L* KD cells. **(H)** Invasion of NCH601 *(sh*Scramble *vs. MTHFD1L* KD) cells into the contralateral hemisphere of NU/NU Nude female mice after 2 months. Each dot represents an individual mouse; mean ± SD; Unpaired *t*-test with Welch’s correction.

**Supplementary Figure 2.**
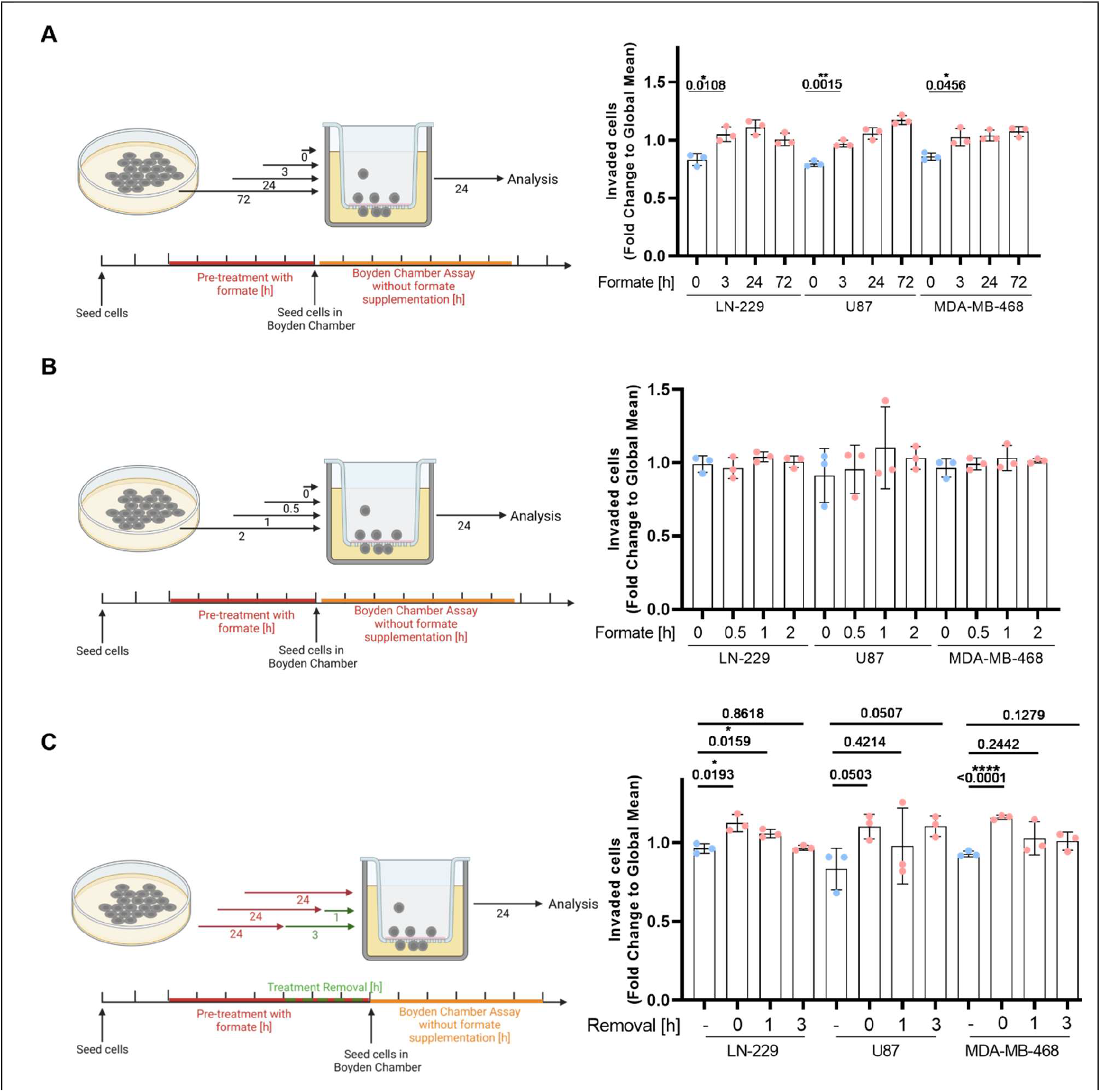
**(A)** On the left: Shematic representation of the experimental layout. On the right: Invasion of LN-229, U87 and MDA-MB-468 cells pre-treated for 0, 3, 24, or 72 hours with 500 μM Na-formate. The invasion was assessed using ECM-collagen-coated Boyden chambers. Each dot represents an independent experiment, the bars represent the mean, and the error bars visualize the standard deviation (SD). The data was evaluated using an unpaired *t*-test with Welch’s correction. **(B)** On the left: Shematic representation of the experimental layout. On the right: Invasion of LN-229, U87, and MDA-MB-468 cells pre-treated for 0, 0.5, 1, or 2 hours with 500 μM Na-formate. The invasion was assessed using ECM-collagen-coated Boyden chambers. Each dot represents an independent experiment, the bars represent the mean, and the error bars visualize the standard deviation (SD). The data was evaluated using an unpaired *t*-test with Welch’s correction. **(C)** On the left: Shematic representation of the experimental layout. On the right: Invasion of LN-229, U87, and MDA-MB-468 cells, which were pre-treated for 24 hours with formate and then cultured for 1 or 3 hours in a neutral medium. The invasion was assessed using ECM-collagen-coated Boyden chambers. Each dot represents an independent experiment, the bars represent the mean, and the error bars visualize the standard deviation (SD). The data was evaluated using an unpaired *t*-test with Welch’s correction.

**Supplementary Figure 3.**
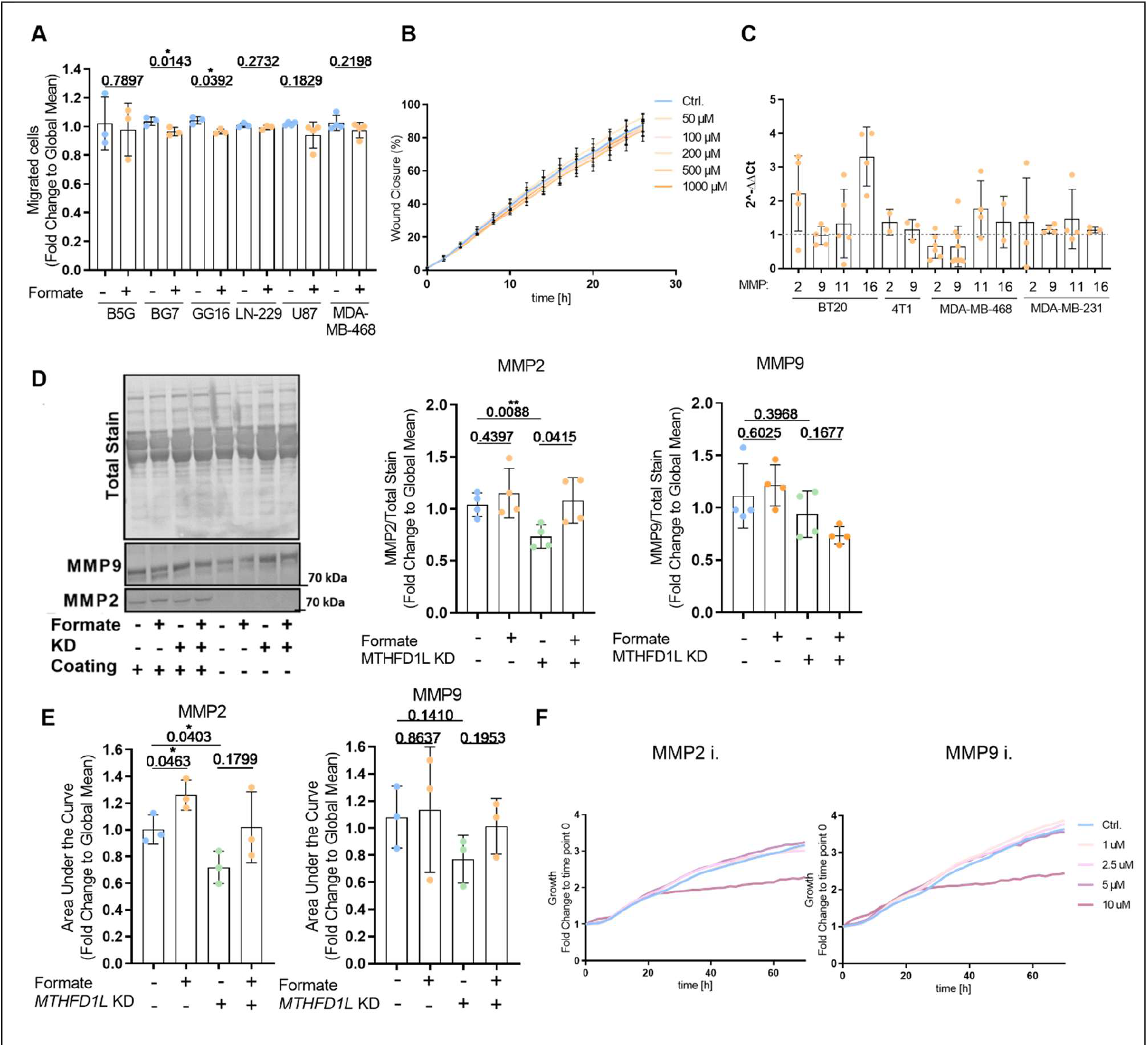
**(A)** Migration of different cells: BG5, BG7, GG16, LN-229, U87, and MDA-MB-468 treated for 24 hours with 500 μM formate was assed using non-coated Boyden chambers. Each dot represents an independent experiment; mean ± SD; Unpaired *t*-test with Welch’s correction. **(B)** Migratory potential of LN-229 cells in response to different formate concentrations (0 μM, 50 μM, 100 μM, 200 μM, 500 μM, and 1mM) as time-dependent quantification of relative wound density in IncuCyte live cell analysis. Graph shows mean ± SEM (*n* = 3). **(C)** mRNA expression from MMP2, MMP9, MMP11, and MMP16 in BT20, 4T1, MDA-MB-468, and MDA-MB-231 cells which were treated for 24 hours with 500 μM formate relative to untreated cells as measured using real-time RT-qPCR. Each dot represents an independent experiment; mean ± SD. **(D)** Expression of MMP2 or MMP9 in the cell culture medium. The total stain served as a loading control. The signal intensity of MMP has been quantified with respect to the signal intensity of the total stain. Each dot represents an independent experiment, the bars represent the mean, and the error bars visualize the standard deviation. The data was evaluated using an unpaired *t*-test with Welch’s correction. **(E)** MMP2 and MMP9 activity of LN-229 *sh*Scramble or *MTHFD1L* KD cells treated for 72 hours with 500 μM formate was assed using gelatin zymography assay. Each dot represents an independent experiment; mean ± SD; Unpaired *t*-test with Welch’s correction. **(F)** Proliferation of LN-229 cells treated for 72 h with different concentrations (0 μM, 1 μM, 2.5 μM, 5 μM, and 10 μM) of MMP2 or MMP9 inhibitor was determined as a time-dependent quantification of cell density (confluence) in IncuCyte. Results are displayed as a fold change to time point zero (*n* = 3).

**Supplementary Figure 4.**
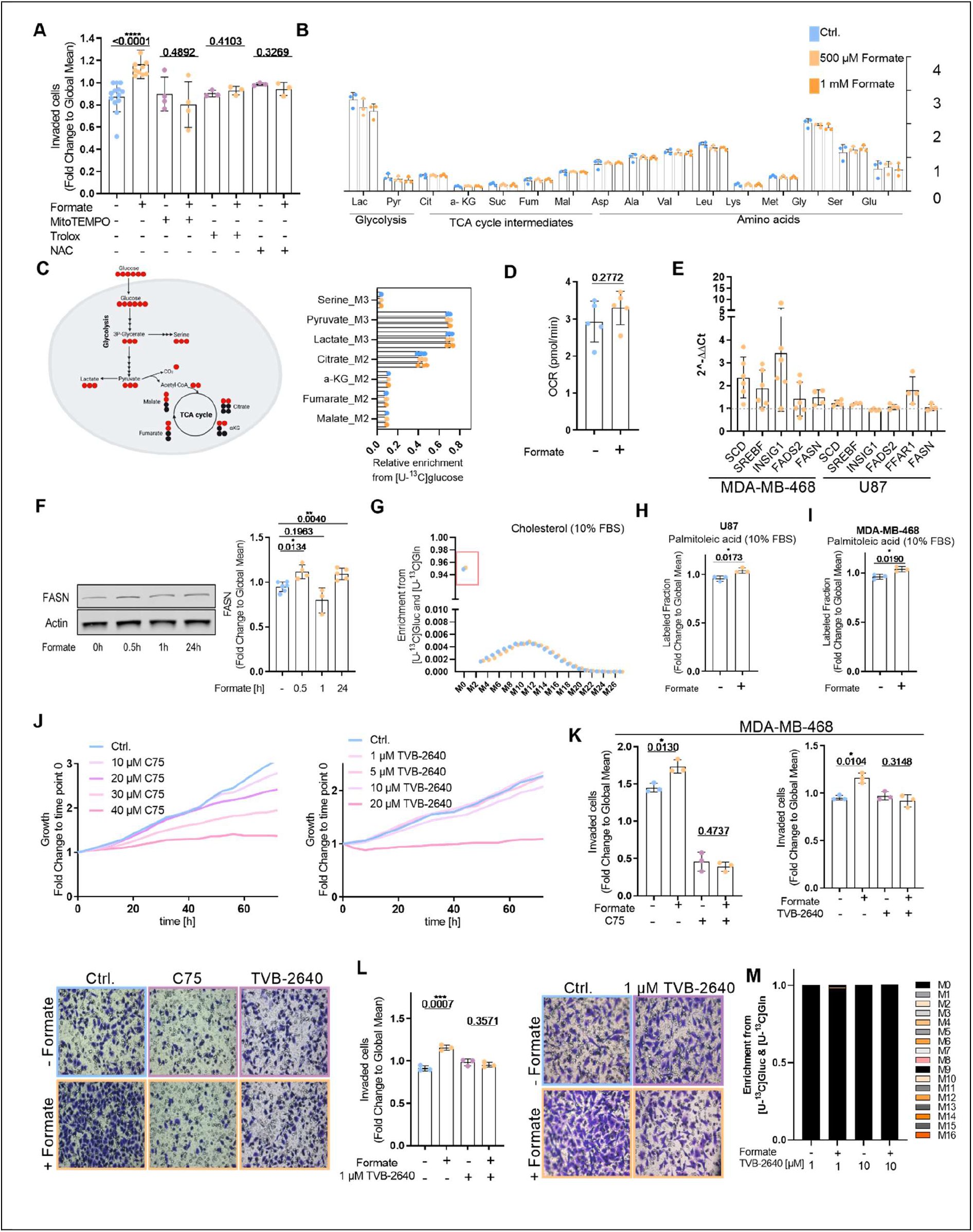
**(A)** Invasion of LN-229 cells treated for 24 h with 500 μM Formate and a Reactive Oxygen Species scavenger (10 μM MitoTEMPO, 5 μM Trolox, or 3 mM NAC) was assed using ECM-collagen coated Boyden chambers. Each dot represents an independent experiment; mean ± SD; Unpaired *t*-test with Welch’s correction. **(B)** Relative Peak Area normalised to packed cell volume (PCV) of different metabolites in LN-229 cells which were treated for 24 h with 500 μM or 1 mM formate. Each dot represents an independent experiment with triplicate wells. **(C)** On the left: Schematic depicting interdependence of glycolysis and TCA cycle as well as expected metabolic ^13^C label pattern from [U-^13^C]glucose tracer. On the right: Enrichment of representative isotopologues of serine, pyruvate, lactate, citrate, α-KG, fumarate and malate upon [U-^13^C]glucose tracer in response to 24 hours, 500 μM or 1 mM formate treatment in LN-229 cells. Each dot represents an independent experiment with triplicate wells. **(D)** Basal cellular respiration in response to 24 h 500 μM formate treatment was determined in LN-229 cells as quantification of mitochondrial oxygen consumption rate (OCR). Each dot represents an individual experiment composed of six technical replicates; mean ± SD. **(E)** mRNA expression from SCD, SREBF, INSIG1, FADS2 and FFAR1 in U87 and MDA-MB-468 cells which were treated for 24 h with 500 μM formate relative to non-treated cells as measured using real-time RT-qPCR. Each dot represents an independent experiment. **(F)** Expression of FASN in LN-229 cells treated with 500 μM formate for different durations (0, 0.5, 1, and 24 hours). The signal intensity of FASN was quantified with respect to the signal intensity of β-actin. Each dot represents an independent experiment, the bars represent the mean, and the error bars visualize the standard deviation (SD). The data was evaluated using an unpaired *t*-test with Welch’s correction. **(G)** Representative graph of isotopologues enrichement of Cholesterol in DMEM supplemeneted with 10% FBS upon [U-^13^C]glucose and [U-^13^C]glutamine tracer response to 72 h 500 μM formate treatment in LN-229 cells. Experiment was repeated 3 times with triplicate wells. **(H)** Labeled fraction of Hexadecenoic acid 2 in DMEM supplemented with 10% FBS upon [U-^13^C]glucose and [U-^13^C]glutamine tracer in response to 72 h 500 μM formate treatment in U87 cells. Each dot represents an independent experiment with triplicate wells; mean ± SD; Unpaired *t*-test with Welch’s correction. **(I)** Labeled fraction of Hexadecenoic acid 2 in DMEM supplemented with 10% FBS upon [U-^13^C]glucose and [U-^13^C]glutamine tracer in response to 72 h 500 μM formate treatment in MDA-MB-468 cells. Each dot represents an independent experiment with triplicate wells; mean ± SD; Unpaired *t*-test with Welch’s correction. **(J)** Proliferation of LN-229 cells treated for 72 h with different concentrations (0 μM, 10 μM, 20 μM, 30 μM, and 40 μM) of C75 and (0 μM, 1 μM, 5 μM, 10 μM, and 20 μM) TVB-2640 was determined as a time-dependent quantification of cell density (confluence) in IncuCyte. Results are displayed as a fold change to time point zero (*n* = 3). **(K)** Invasion of MDA-MB-468 cells treated for 24 h with 500 μM formate and a FASN inhibitor (10 μM C75 and 10 μM TVB-2640) was assed using ECM-collagen coated Boyden chambers. Each dot represents an independent experiment; mean ± SD; Unpaired *t*-test with Welch’s correction. **(L)** Invasion of LN-229 cells treated for 24 h with 500 μM formate and 1 μM TVB-2640 was assed using ECM-Collagen coated boyden chambers. Each dot represents an independent experiment; mean ± SD; Unpaired *t*-test with Welch’s correction. **(M)** Representative graph of isotopologue enrichment of hexadecenoic acid after U-^13^C]glucose and [U-^13^C]glutamine tracer supplementation in response to 72 hours of 1 μM TVB-2640 and 10 μM TVB-2640 treatment in LN-229 cells.

**Supplementary Table 1:**
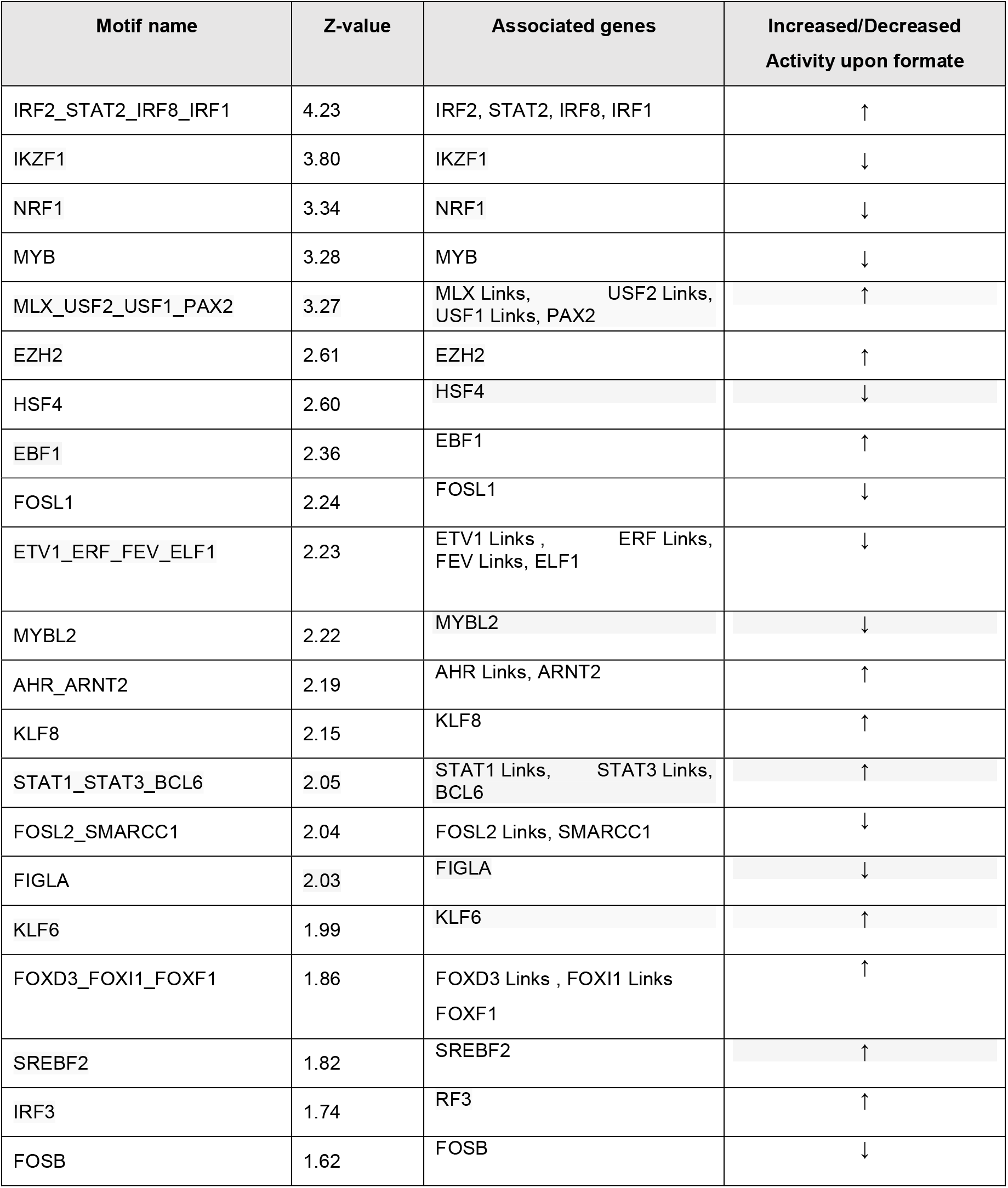
Top Hits of the “Integrated System for Motif Activity Response Analysis” (ISMARA) using the data from the RNAseq (Ctrl versus 24 hours treatment). Represented hits all have a Z-value above 1.5. The arrows indicate if the activity is increased or decreased upon formate treatment.

